# A subset of Haematopoietic Stem Cells resists *Plasmodium* infection-induced stress by uncoupling interferon sensing and metabolic activation

**DOI:** 10.1101/2025.04.09.647965

**Authors:** Christiana Georgiou, Qi Liu, Sara Gonzalez-Anton, Federica Bruno, Flora Birch, Sarah J. Kinston, Shirom Chabra, Nicola K. Wilson, Constandina Pospori, Andrew M. Blagborough, Berthold Göttgens, Tiago C. Luis, Cristina Lo Celso

## Abstract

Hematopoietic stem cells (HSCs) sustain lifelong haematopoiesis as their progeny differentiate into all blood cell lineages. Homeostatic HSCs are mostly quiescent and only rarely divide, however their proliferation and differentiation rates can be modulated by external factors. Acute and chronic infections from a wide range of pathogens are known to challenge HSCs at the population level, being forced to respond to inflammation-mediated organismal demand to replenish the myeloid cell pool. However, less is known about the degree of heterogeneity in the HSCs’ response to inflammation at the single cell level. Here, using a natural murine malaria model and an NHS-ester biotin dilution assay we identify two subsets of HSCs, Biotin^Lo^ and Biotin^Hi^, with distinct proliferation kinetics. Using combined functional, single-cell transcriptomics and phenotypic analyses, we uncover that Biotin^Hi^ HSCs remain highly functional despite expressing strong interferon response signatures. These infection-resistant HSCs express high levels of MHC II and are metabolically distinct from the remaining HSCs as they maintain less active mitochondria. These findings demonstrate that a likely reserve pool of HSCs remains highly functional during *Plasmodium* infection not because cells are shielded, but because they maintain a stemness associated metabolic profile despite effectively sensing inflammation.

## INTRODUCTION

Hematopoiesis is the continuous process through which hematopoietic stem cells (HSCs) sustain the production of all blood cells, ensuring oxygen transportation, clotting, and immune surveillance and responses. The hematopoietic system is regulated to maintain homeostasis, with HSCs balancing their quiescence and proliferation, and self-renewal and differentiation throughout life to fulfil physiological demands. Severe infections are a strong cell-extrinsic stress because they cause strong inflammatory signals that activate HSCs to proliferate and replenish the downstream progenitor pool. Following several studies involving pathogen-mediated infection (Haltalli et al., 2020; Matatall et al., 2016) or sterile inflammation models (Essers et al., 2009; Yamashita, Nitta and Suda, 2015; Pietras et al., 2016; Bogeska et al., 2022), it is widely agreed that continuous exposure to proliferation-induced stress leads to HSC loss and functional decline (Essers et al., 2009; Baldridge et al., 2010; Hirche et al., 2017; Bogeska et al., 2022). However, it would be evolutionarily disadvantageous if external stresses cause attrition of the entire HSC pool, and we thus hypothesized that the HSC response to infection ought to be heterogeneous and at least a subpopulation of functional HSCs would withstand infection-derived stress.

Malaria is a life-threatening disease with pathology caused by *Plasmodium* parasites invading erythrocytes. Recently, more than 263 million clinical infections and 597,000 malaria deaths were reported annually (World Malaria Report 2024, 2024), placing malaria among the top lethal infectious diseases in humans, along with HIV and tuberculosis (Coban et al., 2018). Malaria cases are on the rise globally, driven by climate change (Samarasekera, 2023) and emerging resistance to front-line antimalarials (Ashley et al., 2018; Venugopal et al., 2020), and it is important that we better understand how HSCs are affected by the strong inflammation triggered by *Plasmodium* infection and what mechanisms may be in place to withstand it. Murine malaria models have started to shed light on the complexity of malaria infection and its detrimental effects on hematopoietic cells and stem cells. Using the murine parasite *Plasmodium berghei* we previously demonstrated hematopoietic perturbations in hematopoietic progenitors, especially the expansion of Sca-1^+^ progenitors, in host animals infected through the bites of sporozoite-positive *P. berghei* infected mosquitos (Vainieri et al., 2016; Haltalli et al., 2020). In addition, we reported loss of HSC transcriptional identity and function, linked to loss of osteoblasts, increased bone marrow vasculature leakiness and increased HSC Reactive Oxygen Species (ROS) in HSCs (Haltalli et al., 2020).

Here, we address the question of heterogeneity in the HSC response to malaria infection-driven inflammation. Using the gold standard HSC assay of serial transplantation, we ask whether a subset of rare, quiescent HSCs may maintain stemness, despite malaria infection. Specifically, we assess whether malaria infection induces proliferation of all HSCs equally, and what mechanisms may underpin HSCs’ heterogeneous responses. Through a combination of molecular, phenotypic and functional analyses we not only identify a quiescent subset of HSCs that preserves function throughout malarial infection, but also reveal possible determinants of these infection-resistant HSCs.

## RESULTS

### Serial transplantation reveals residual HSC function despite severe acute infection

To identify heterogeneities in the response of HSCs to the inflammation resulting from a natural infection we developed an infection set-up based on the rodent malaria parasite *P. berghei* ANKA 2.34, which we had previously shown to cause extensive functional damage to HSCs (Haltalli et al., 2020). As the liver stage of sporozoite-mediated *Plasmodium berghei* infection does not affect hematopoietic stem and progenitor cells (Haltalli et al., 2020), here we used a blood-stage infection model, which reliably leads to parasitaemia >3% in 5 rather than 7 days from infection (Figure 1A and B). In brief, experimental mice are infected with parasitized blood (P1) obtained by injecting primary donors with aliquots of frozen blood stocks (P0) harvested from sporozoite-infected donors. Experimental mice were analyzed at day 5 post blood infection (pbi),before cerebral malaria signs develop (Figure 1A). Flow cytometry analysis of bone marrow (BM) from control and infected animals confirmed the hallmark expansion of LKS (Lin^neg^c-Kit^+^SCA-1^+^) compartment, containing multipotent progenitors (MPPs) and HSCs (Figure 1C), consistent with the sporozoite infection model (Haltalli et al., 2020). To identify HSC heterogeneity in the context of malaria, we focused our studies on as CD48^neg^ HSCs (LKS CD150^+^ CD48^neg^ cells), a small subpopulation of SLAM HSPCs highly enriched for functional HSCs (Akinduro et al., 2018). The absolute number of these cells was dramatically reduced in infected mice (Figure 1D), and because we previously showed protection of HSC function in settings where CD48^neg^ HSCs were maintained (Haltalli et al., 2020), we therefore set out to test whether any functionality may be retained within the CD48^neg^ HSC compartment that naturally survives during the infection. We sort-purified CD48^neg^ HSCs from mTmG transgenic (Muzumdar et al., 2007)control and infected donor mice, injected 120 tomato+ CD48^neg^ HSCs into each conditioned CD45.1 recipient together with support CD45.1 whole bone marrow (WBM) cells (Figure S1A), and assessed their reconstitution ability by measuring tomato+ donor-derived peripheral blood (PB) cells in all recipients through flow cytometry analyses (Figure 1E and Figure S1B). Cells from infected donors generated fewer progeny compared to control, consistent with previous infection/inflammation models (Haltalli et al., 2020; Baldridge et al., 2010; Matatall et al., 2016; Bogeska et al., 2022); however, their output was consistently evident throughout the assay (Figure 1E), with no indication of any myeloid or lymphoid lineage bias compared to the output of control CD48^neg^ HSCs (Figure S1B). As a reduced, but still detectable BM reconstitution was present in the recipients of infected donors’ HSCs (Figure 1F), we tested the self-renewal ability of the injected HSCs by performing secondary WBM transplants (Figure S1C). Once normalised for primary transplant output, HSCs from infected donors showed secondary reconstitution ability equivalent to that of controls in the PB (Figure 1G), and a more variable but still overall equivalent reconstitution ability in BM (Figure 1H). These data suggested that while severe infection does lead to a reduction in HSC functionality, this is not completely lost, raising the question of whether a HSC subpopulation may exist that can withstand infection-driven inflammation.

**Figure 1.**
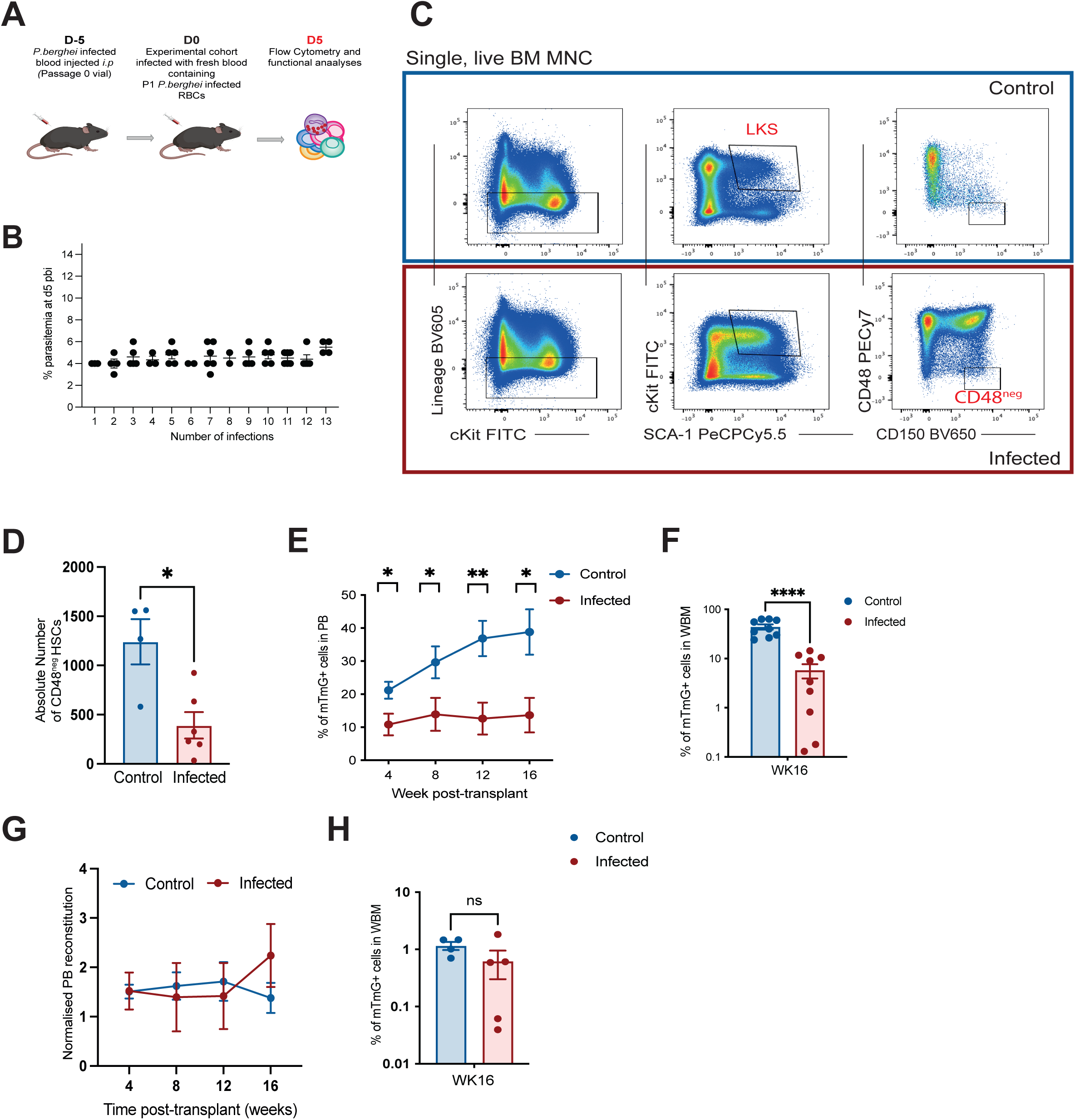
Detectable HSC function in mice undergoing severe infection. (A) Experimental set-up for infected cohort. (B) Parasitemia at day 5 post blood infection. Each dot represents a mouse, data collected from 13 independent infections (C) FACS gating strategy for the identification of CD48^neg^ HSCs in c-Kit enriched BM MNCs from control and infected mice. Plots are representative of 65 independent infection experiments. (D) Absolute number of CD48^neg^ HSCs in control and infected animals, n= 4 and 6 infected animals from 2 independent infections. (E) Peripheral blood (PB) engraftment in primary transplant recipients of 120 mTmG+ CD48^neg^ HSCs from control and infected mice, measured over 16 weeks. (F) Percentage of mTmG+ cells in WBM of primary transplant recipients. For e and f n= 9 and 9 recipients of HSCs from control and infected donors, pooled from 2 independent transplantation experiments. (G) PB reconstitution of secondary transplant recipients normalised to primary transplant PB reconstitution at 16 weeks. Engraftment was measured as the percentage of mTmG+ cells in PB. (H) Percentage of mTmG+ cells in WBM of secondary transplant recipients. In G and H n= 4 and 5 recipients of WBM from primary recipients in control and infected groups, pooled from 2 independent experiments. In B and D-H each dot represents one animal. Error bars represent mean ±s.e.m. Statistically significant differences were determined using Student’s t-test and for multiple comparisons one way ANOVA with post-Tukey correction (**** represents p≤ 0.0001; *** represents p≤0.001; ** represents p≤0.01; * represents p≤0.05).

### A primitive HSC subset does not proliferate despite malaria-induced stress

To investigate HSC heterogeneous responses to *P. berghei* infection, we employed the previously described N-hydroxysulfosuccinimide (NHS)-ester-LC-LC-biotin (NHS-ester-biotin from now on) labelling dilution assay (Luis TC et al., 2023; Nygren and Bryder, 2008), which constitutes a non-invasive method to investigate the proliferation history of HSPCs while allowing the isolation of subsets with distinct proliferation kinetics for functional analysis, without the need of complex transgenic systems. We included MPP populations (MPP2, LKS Flt3^neg^ CD150^+^ CD48^+^, MPP3, LKS Flk2^neg^ CD150^neg^ CD48^+^; MPP4, LKS Flk2^+^ CD150^neg^ CD48^+^) in our analysis as an internal control for cells that divide rapidly during *Plasmodium* infection and included EPCR as a marker of highly functional HSCs (Kent et al., 2009). We therefore analyzed in parallel the CD48^neg^ HSC population and its EPCR+ fraction, referred to as E-SLAM HSCs from now on (Figure S2A). As an initial control, we confirmed that all primitive HSPC populations were effectively and homogenously labelled by a single injection of the NHS-ester-biotin (Figure S2B).

To identify heterogeneity in HSC proliferation during infection, we injected the NHS-ester-biotin intravenously (*i.v.*) in control and day 3 pbi infected animals and performed subsequent analyses at day 5 pbi (Figure 2A). As expected, MPP populations in infected mice showed considerable biotin dilution, with MPP2 presenting the strongest and MPP4 the weakest reduction in NHS-ester-biotin MFI (Figure S2C), consistent with previously described emergency myelopoiesis in this system (Haltalli et al., 2020; Belyaev et al., 2013). Within the HSC compartment, CD48^neg^ HSCs and E-SLAM HSCs in control animals retained higher NHS-ester-biotin levels and 92% CD48^neg^ and 93% E-SLAM cells remained Biotin^Hi^, consistent with their well-documented quiescence. Instead, in infected animals NHS-ester-biotin dilution was evident as the biotin label MFI of both the overall populations was significantly reduced (Figure 2B and C). As a result, we could split HSC populations according to NHS-ester-biotin levels in infected animals as Biotin^Hi^ cells that retained biotin label (43% CD48^neg^ and 45% E-SLAM Biotin^Hi^), and Biotin^Lo^ cells that had diluted NHS-ester-biotin, suggesting increased proliferation in this population (Figure 2B, 2C and Figure S2D).

**Figure 2.**
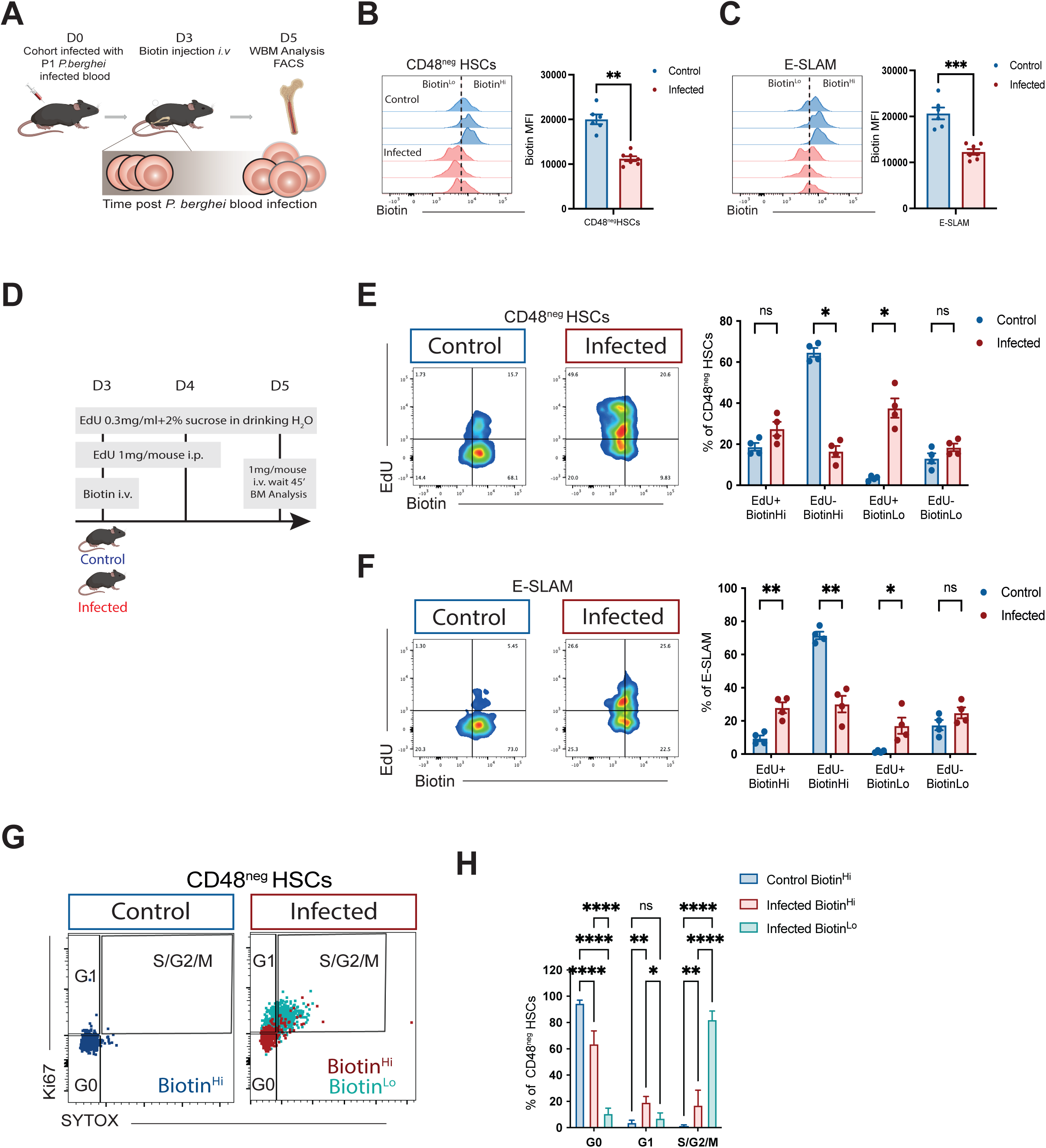
Biotin dilution identifies HSC subpopulation that does not proliferate in infected mice. (A) Experimental set-up for the measurement of NHS-ester-biotin label dilution during infection. Mice received the label at day 3 pbi. (B) Representative histograms and quantification of NHS-ester-biotin mean fluorescence intensity (MFI) on CD48^neg^ HSCs from control (blue) and infected (red) mice. Dotted line represents the threshold between Biotin^Hi^ and Biotin^Lo^ cells. (C) Representative histograms and quantification of NHS-ester-biotin mean fluorescence intensity (MFI) on E-SLAM HSCs. In B and C n=6 and 7 control and infected mice, respectively, pooled from 2 independent infections. (D). Experimental set up with chronic EdU administration to identify HSCs that proliferate during infection. (E) Representative flow cytometry dot plots showing EdU uptake and NHS-ester-biotin dilution in CD48^neg^ HSCs from control and infected mice (left), indicating that the majority of CD48^neg^ HSCs are Biotin^Hi^ and EdU negative in control mice, while a higher proportion of cells are EdU+, biotin^Lo^ or both in infected mice. (F) Representative dot plots and quantification of EdU incorporation and NHS-ester-biotin dilution in E-SLAM HSCs. In E and F n= 4 control and 4 infected mice. (G) Representative dot plots of Ki67 immunostaining and DNA content (SYTOX label) in Biotin^Hi^ CD48^neg^ HSCs from control mice (left) and in Biotin^Hi^ and Biotin^Lo^ CD48^neg^ HSCs from infected mice (right). (H) Proportion of Biotin subsets of CD48^neg^ HSCs from control (n=4) and infected mice (n=3). Each dot represents one animal. Error bars represent mean ±s.e.m in B, E and F, and s.d. in H. Statistically significant differences were determined using Student’s *t*-test or two-way ANOVA with post-Tukey correction for multiple comparisons (**** represent p≤ 0.0001; *** represent p≤0.001; ** represent p≤0.01; *represent p≤0.05).

To validate that the reduction of NHS-ester-biotin labelling in HSCs was due to proliferation-induced label dilution, we chronically administered EdU to infected and control mice (Figure 2D). As expected, in control mice the majority of CD48^neg^ HSCs were Biotin^Hi^ and EdU^neg^. Instead, in infected mice the majority of CD48^neg^ HSCs were Biotin^Lo^ and EdU^+^ (Figure 2E). Consistent with EPCR marking very primitive, quiescent HSCs (Kent et al., 2009), in control animals fewer E-SLAM HSCs than CD48^neg^ HSCs were EdU^+^. This was the case also in infected animals, where E-SLAM HSCs were uniformly split between Biotin^Hi^ and Biotin^Lo^, EdU^+^ and EdU^neg^ (Figure 2F). The high increase in EdU^+^ Biotin^Lo^ HSC fractions suggested that indeed NHS-ester-biotin was diluted as a result of proliferation. To further validate our finding, we performed cell cycle analysis on Biotin^Hi^ and Biotin^Lo^ HSC populations. Consistent with the EdU incorporation data, in control animals, most CD48^neg^ Biotin^Hi^ HSCs were in G0, and in infected animals the largest portion of CD48^neg^ Biotin^Hi^ HSCs were enriched in G0, while CD48^neg^ Biotin^Lo^ cells were mostly in S/G2/M phases of the cell cycle (Figure 2G and 2H). Together, these data indicated that the proliferative response of HSCs to infection is heterogeneous, and that a fraction of HSCs does not proliferate despite a highly inflammatory *milieu*.

### E-SLAM Biotin^Hi^ cells retain self-renewal and multipotency

To test whether Biotin^Hi^ HSCs remain functional despite *Plasmodium* infection-driven stress and how they would compare to Biotin^Lo^ HSCs from infected mice and, importantly, Biotin^Hi^ HSCs from control healthy animals, we set up serial transplantation assays. We sort-purified Biotin^Hi^ and Biotin^Lo^ E-SLAM HSCs from CD45.2 infected mice and Biotin^Hi^ E-SLAM HSCs from CD45.2 control mice and injected 10 cells from one of the three populations in CD45.1 lethally irradiated recipients, together with 250,000 CD45.1 WBM support cells (Figure 3A). We did not assess the functionality of Biotin^Lo^ E-SLAM HSCs from control mice because they were too rare. Longitudinal analysis of peripheral blood reconstitution in the three recipient groups indicated a surprising trend for Biotin^Hi^ cells from infected donors to achieve the highest levels of engraftment, while Biotin^Lo^ cells achieved lower engraftment (Figure 3B and 3C). The lineage differentiation potential of the three HSC populations was also similar, with Biotin^Lo^ cells showing a trend for higher myeloid differentiation at earlier time points (Figure 3D), consistent with emergency granulopoiesis being triggered in the donor mice (Haltalli et al., 2020). Whole BM reconstitution, assessed at the end of the assay, was variable and with a similar average in all three groups, though a higher proportion of recipients of Biotin^Hi^ E-SLAM HSCs from infected donors showed high engraftment (>25% engraftment in 5/11 Biotin^Hi^ control, 9/11 Biotin^Hi^ infected and 2/4 Biotin^Lo^ infected; Figure 3E). Consistent with this, the average frequency of donor-derived HSCs was similar across all three recipients’ groups; however, 2/11 Biotin^Hi^ control, 5/11 Biotin^Hi^ infected and 1/4 Biotin^Lo^ infected recipients had reconstitution >25% (Figure 3F). Next, we injected 2 x 10^6^ WBM cells from highly reconstituted primary recipients from each initial HSC population into lethally irradiated CD45.1 secondary recipients (Figure 3G). All secondary recipients showed donor-derived engraftment in the blood, however the mice from the E-SLAM Biotin^Hi^ infected donors group had significantly higher engraftment than those from the E-SLAM Biotin^Hi^ control and E-SLAM Biotin^Lo^ infected groups (Figure 3H). The lineage output of the three E-SLAM HSC subpopulations in secondary recipients was similar (Figure 3I). BM analysis at the end of the assay confirmed that also in this compartment E-SLAM Biotin^Hi^ cells from infected donors conferred significantly higher reconstitution compared to either E-SLAM Biotin^Hi^ control and –Biotin^Lo^ infected cells (Figure 3J), and repopulation of the HSC compartment was similar across groups (Figure 3K). These data indicated that NHS-ester-biotin label retention in the context of severe infection was linked with the highest repopulation potential, and therefore identified infection-resistant HSCs (IR-HSCs).

**Figure 3.**
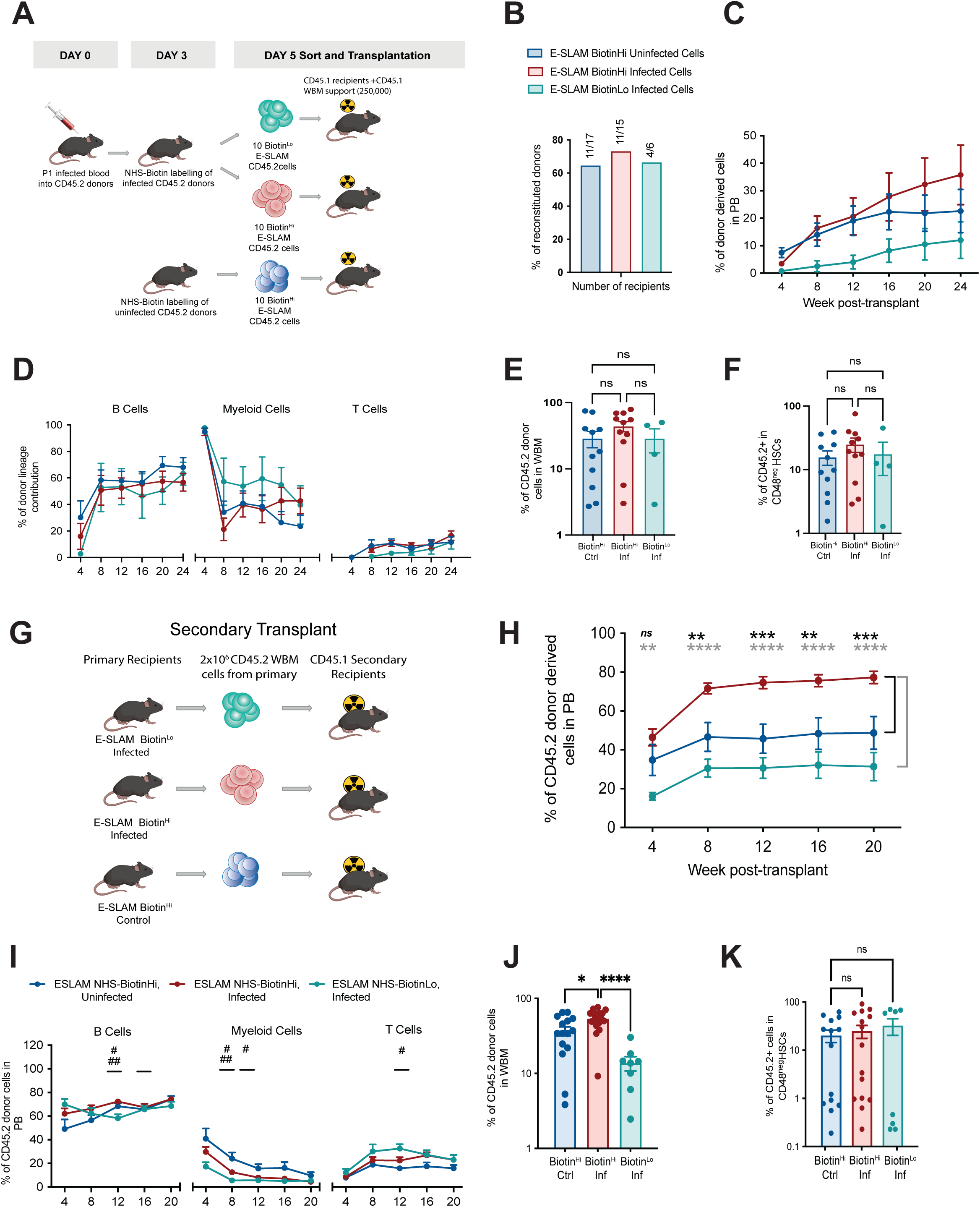
Biotin^Hi^ HSCs from infected mice retain highly engraftment potential. (A) Experimental set up for primary transplants. 10 CD45.2 E-SLAM Biotin subset cells per condition were injected into lethally irradiated CD45.1 recipients together with 250,000 BM MNCs from CD45.1 donors as support. (B) Percentage of reconstituted CD45.1 recipients, ie mice with detectable CD45.2 cells in PB. Numbers above each bar indicate the number of mice reconstituted out of total animals injected. (C and D) Overall PB reconstitution and multilineage output measured in reconstituted recipients of cells from each subset. (E and F) BM and CD48^neg^ HSC compartment reconstitution in recipients of cells from each subset, measured at 24 weeks post-transplant. In C to F n= 11, 11 and 4 recipients of Biotin^Hi^ control, Biotin^Hi^ infected and Biotin^Lo^ infected HSC subsets, pooled from two independent transplantation experiments. (G) Experimental set up. Mice received 2 million WBM MNCs from primary recipients per condition. Donors were primary recipients with BM engraftment >20%. (H and I) Overall PB reconstitution and multilineage output in secondary recipients. (J and K) BM and CD48^neg^ HSC compartment reconstitution in secondary recipients at 20 weeks post-transplant. In H to K n= 15, 17 and 8 recipients of BM from primary recipients of Biotin^Hi^ control, Biotin^Hi^ infected and Biotin^Lo^ infected HSC subsets, pooled from two independent experiments. The data are presented as means ± s.e.m. The p values were determined by Student’s *t* test or two-way ANOVA with post-hoc Bonferroni correction for multiple comparisons. ****p ≤ 0.0001;** (or ##) p ≤ 0.01; * p<0 (or #). # Shows significance between different groups (## in B cells between Biotin^Lo^ and Biotin^Hi^ Infected and # between Biotin^Lo^ infected vs Biotin^Hi^ control at the time points indicated). In C, D and I no statistically significant differences between groups were identified.

### Infection-resistant HSCs maintain gene signatures associated with stem cell function

To understand and highlight the molecular changes taking place in Biotin HSC subsets as a consequence of *P. berghei* infection, we index sort-purified CD48^neg^ HSCs from 8 control and 7 malaria burdened mice. All cells were processed for single cell RNA-sequencing (scRNA-seq) and we retrospectively assigned them to HSC subsets based on biotin label intensity from index-sorting data (Figure 4A-B). 58 Biotin^Hi^ HSCs from control and 64 from infected mice, respectively, and 43 Biotin^Lo^ HSCs from infected mice were retained after quality control, for a total of 165 cells with an average of 8026 genes detected per cell (Figure 4A). Biotin^Hi^ CD48^neg^ HSCs expressed higher levels of EPCR than Biotin^Lo^ cells (Figure 4B, right), which was consistent with E-SLAM HSCs being enriched for Biotin^Hi^ cells (Figure 2C) and with Biotin^Hi^ cells being more functional in serial reconstitution assays than Biotin^Lo^ cells (Figure 3). First, we identified 2,163 highly variable genes (HVGs) across the three populations and created a uniform manifold approximation and projection (UMAP) space to capture nuanced relationships between HSCs in each of the 3 groups (Biotin^Hi^ control/infected, Biotin^Lo^ infected). Biotin^Hi^ and Biotin^Lo^ HSCs from infected mice appeared quite interspersed with each other, whereas the majority of the control cells were separated from the infected condition ones (Figure 4C).

**Figure 4.**
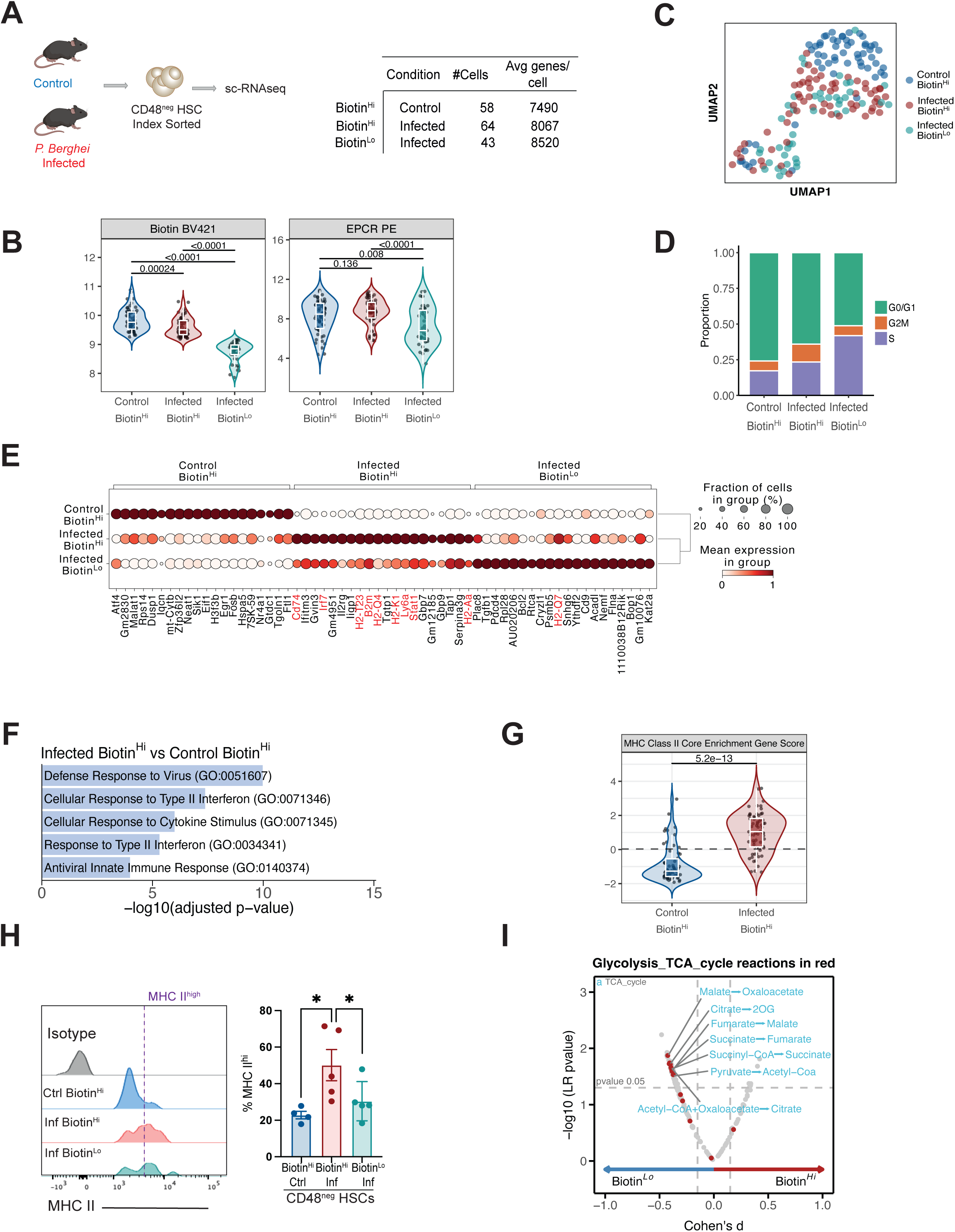
Infection-resistant HSCs have a unique transcriptional profile. (A)Experimental set up. CD48^neg^ HSCs were index-sorted from control (n = 7) and infected (n = 7) mice from two independent infections and processed for scRNA sequencing. (B) Index sort values for biotin label (left) and EPCR expression (right). (C) UMAP analysis shows Biotin^Hi^ HSCs from control animals clustering separately and Biotin^Hi^ and Biotin^Lo^ HSCs from infected mice mixing together. (D) Cell cycle related signatures show Biotin^Hi^ HSCs from infected mice to have an intermediate profile. (E) Dotplot highlighting top 15 DEGs in each cell category, and confirming Biotin^Hi^ HSCs from infected mice being more similar to Biotin^Lo^ HSCs from infected mice than Biotin^Hi^ HSCs from control mice. Genes in red are found in the GO categories ‘Cellular response to type II interferon’ and ‘Response to type II interferon’ shown in Fig. 4F. (F) Results of Gene Ontology analysis for DEGs between control and infected Biotin^Hi^ HSCs. (G) MHC score analysis highlights higher expression of MHC class II signatures (cd74, H2-Aa, Hs-Eb1, H2-Ab1, Ctss and Tubb) in Biotin^Hi^ HSCs from infected mice compared to control. (H) Representative Flow cytometry analysis of MHC II expression in CD48^neg^ HSCs. The data are presented as means ± s.e.m. n= 8 control and 9 infected mice from two independent infections. The p values were determined by unpaired Student’s *t*-test or ordinary one-way ANOVA with Tukey correction. * p<0 05. (I) Biosynthetic pathways analysis. In B and G P values were determined by Wilcoxon test.

Next, we calculated the putative cell cycle state of the different HSC populations, and could see that whilst the majority of the cells were calculated to be in G0/G1 phase, the number of cells in S phase was higher in Biotin^Lo^ cells (Kowalczyk et al., 2015; Nestorowa et al., 2016) (Figure 4D). The majority of Biotin^Hi^ HSCs from control mice expressed high levels of *Egr1, Fosb*, which is consistent with a G0/G1 transcriptional signature, since these genes specifically inhibit proliferation, and have been shown previously to regulate homeostasis of the HSC compartment (Min et al., 2008). Furthermore, the G0/G1 signature of Biotin^Hi^ HSCs is supported by their low expression of genes associated with cell cycle activation (*Cd69*) (Scheicher et al., 2015) and progression (*Cdk6*) (Bujanover et al., 2020; Laurenti et al., 2015). Conversely, a higher proportion of Biotin^Lo^ cells exhibited a gene expression pattern that reflect cells in S and G2/M cell cycle phases, including low levels of quiescence-associated genes (*Egr1, Fosb*) and high levels of cell cycle activation genes (*Cd69, Cdk6*). Interestingly, Biotin^Hi^ cells from infected mice presented an intermediate pattern of distribution across phases of the cell cycle (Figure 4D), suggesting that these cells may be activated to some degree, but were less likely to be proliferative than Biotin^Lo^ HSCs. These findings were consistent with our flow cytometry analysis of cell cycle distribution of Biotin^Hi^ and Biotin^Lo^ HSCs from control and infected mice (Figure 2G and 2H). The results of UMAP and cell cycle analyses raised the question of whether Biotin^Hi^ HSCs from infected mice may remain functional because of an ability to retain stemness despite being activated by infection.

To try to identify the molecular regulators of the retained “stemness” in the Biotin^Hi^ cells, we utilized a number of previously published gene lists that assess HSC stemness, function and attrition at the transcriptional level. Application of the HSC Score algorithm (Wilson et al., 2015; Hamey and Göttgens, 2019) indicated that Biotin^Hi^ cells expressed transcriptional signatures linked to HSC stemness at significantly higher levels than Biotin^Lo^ cells, and with no significant differences between Biotin^Hi^ cells from control and infected mice (Figure S3A top left panel). Moreover, a similar analysis using a more recently developed Repopulation Score (RepopSig), based on a signature of 23 related genes that enrich for highly engraftable HSCs (Che et al., 2022), generated equivalent results (Figure S3A top right panel), consistent with a model whereby Biotin^Hi^ infected cells are highly functional compared to Biotin^Lo^ cells and similar to Biotin^Hi^ control cells. As proliferation is known to lead to endogenous DNA damage and to promote HSC accelerated ageing associated with p53 activation, upregulation of p53 target genes, and myeloid-biased differentiation (Wang et al., 2023), we asked whether we could identify signs of proliferation-induced stress in HSCs from infected mice. To this end, we computed both the p53 Score (Wang et al., 2023) and the Aging Score (Bogeska et al., 2022) for all cells in the 3 subsets. Biotin^Hi^ cells from control and infected mice presented equivalent values for both scores, while Biotin^Lo^ cells had higher scores but not significantly different to both control and infected groups of Biotin^Hi^ cells (Figure S3A). These data suggest that during *P. berghei* infection Biotin^Hi^ HSCs enter the cell cycle but remain resistant to inflammation-driven attrition, while Biotin^Lo^ HSCs may be primed with an ageing phenotype. The Biotin^Lo^ HSCs’ transcriptional profile is consistent with a previous study where HSCs displayed strong ageing signature and functional attrition after repeated induction of interferon responses (Bogeska et al., 2022).

### Interferon and metabolic transcriptional signatures are uncoupled in Biotin^Hi^ HSCs from infected mice

To characterize the transcriptional profiles associated with the heterogeneous responses of HSCs to malaria infection, we interrogated the scRNA-seq dataset to determine differentially expressed genes (DEGs) that would uniquely characterize the three HSC populations. When we normalised gene expression levels across all three populations (Supplementary Table 1), we confirmed that overall Biotin^Hi^ cells from infected mice clustered more closely with Biotin^Lo^ cells from infected mice (Figure 4E and S3B, top panel). This was because both subsets of HSCs from infected mice expressed strong interferon response signatures, albeit with several unique traits. Specifically, we found that *Cd74* was significantly higher in infected Biotin^Hi^ cells along with other genes associated with HSC maintenance (*Il2rg* and *Stat1*)(Li et al., 2022), transcriptional response to type I interferon (*Irf7*) (Ning et al., 2011), and negative regulation of IFNγ signalling (*Irgm1*) (King et al., 2011) (Figure 4E and Supplementary Table 1). Consistent with this, gene ontology (GO) pathway enrichment analysis using the list of 107 genes upregulated in Biotin^Hi^ infected vs control cells (Figure S3C, left panel and Supplementary Table 2) identified 5 GO terms relating to defence response to virus, and cellular response to cytokines and type II interferon (Figure 4F).

We decided to focus our studies on *Cd74* as it is known to be essential for the trafficking and assembly of MHC class II molecules, hence for antigen presentation, and has been implicated in safeguarding a quiescent subpopulation of HSCs expressing MHC II from stress-induced proliferation (Li et al., 2022). Because *Cd74* expression is associated with MHC class II, we hypothesized that Biotin^Hi^ HSCs from infected mice may express higher levels of MHC II. We developed an MHC II score based on the 6 MHC class II core enrichment genes (*Cd74, H2-Aa, H2-Eb1, H2-Ab1, Ctss, Tubb1*) and indeed Biotin^Hi^ HSCs from infected animals had significantly higher MHC II scores (Figure 4G) compared to Biotin^Hi^ HSCs from control animals. Next, we validated MHC II expression in HSCs at the protein level using flow cytometry. In control animals, a proportion of Biotin^Hi^ HSCs expressed high levels of cell surface MHC-II protein (Figure 4H and Figure S3D). Upon infection, Biotin^Hi^ CD48^neg^ HSCs and E-SLAM HSCs had the highest proportion of cells expressing high levels of MHC II proteins (Figure 4H and Figure S3D, respectively). Interestingly, a high proportion of MPPs already expressed high levels of MHC II in steady state, and only the proportion of MHC II highly expressing MPP4 cells increased significantly upon infection (Figure S3E). These analyses indicated that in infected animals the strongest upregulation of MHC II took place in HSCs, and in particular in Biotin^Hi^ HSCs. The co-existence of stemness and interferon response signatures is therefore a unique characteristic of Biotin^Hi^ HSCs in infected animals, setting them aside from Biotin^Hi^ HSCs from control animals, and raising the question of what mechanisms may support the higher functionality of Biotin^Hi^ compared to Biotin^Lo^ HSCs.

When we compared infected Biotin^Hi^ and Biotin^Low^ cells, we identified 9 DEGs (Figure S3C, right panel), in agreement with our initial UMAP analysis showing a high overlap between all cells from infected animals. Higher *Procr* expression in Biotin^Hi^ cells was consistent with index sorting data (Figure 4B). Higher *Bcl2* expression in Biotin^Low^ cells may explain their relatively high functionality in primary transplants (Figure 3C-E). *Cryzl1* upregulation in Biotin^Low^ cells caught our attention because this gene encodes for a protein that binds NAD(P)H and has high similarity to a quinone oxidoreductase that regulates cell fate decisions in response to stress (Thapa et al., 2020). Given the acknowledged link between metabolic status and HSC functionality, we reasoned that functional differences between the two Biotin cell subsets might be due to metabolic changes taking place in the mitochondria. We therefore took advantage of the deep neural network model scFEA (Alghamdi et al., 2021) and performed computational inference of the biosynthetic activities across all HSC subsets in our scRNA-seq data. The analysis revealed five biosynthetic pathways associated with TCA cycle in Biotin^Lo^ cells from infected animals (Figure 4I) suggesting that Biotin^Lo^ cells are more metabolically active compared to Biotin^Hi^ infected cells, likely linked to their proliferative state.

### Biotin^Hi^ HSCs maintain low mitochondria activity while Biotin^Lo^ HSCs are metabolically primed

Quiescent HSCs are characterized by low oxidative phosphorylation (OXPHOs) and reactive oxygen species (ROS), and are known to highly depend on glycolysis (Papa, Djedaini and Hoffman, 2019; Hinge et al., 2020; Liang et al., 2020; Fernández-Vizarra et al., 2022). To validate whether Biotin^Hi^ HSCs from control and infected animals maintain low metabolic profiles compared to Biotin^Lo^ HSCs we used flow cytometry analyses to measure mitochondrial superoxide (MitoSOX) production, mitochondrial membrane potential (MMP) and mitochondrial mass as an indication of active OXPHOS. We observed that the majority of Biotin^Hi^ HSCs in control and infected animals were not stained by the MitoSOX dye compared to Biotin^Lo^ cells, a high proportion of which was MitoSOX^+^ (Figure 5A and B). Interestingly MitoSOX levels in MPP2, MPP3 and MPP4 were higher in steady state, consistent with these cells being more proliferative than HSCs, but in infected mice a smaller proportion of MPPs was MitoSox positive, and MitoSox levels did not change dramatically as NHS-ester biotin was diluted (Figure S4A).

**Figure 5.**
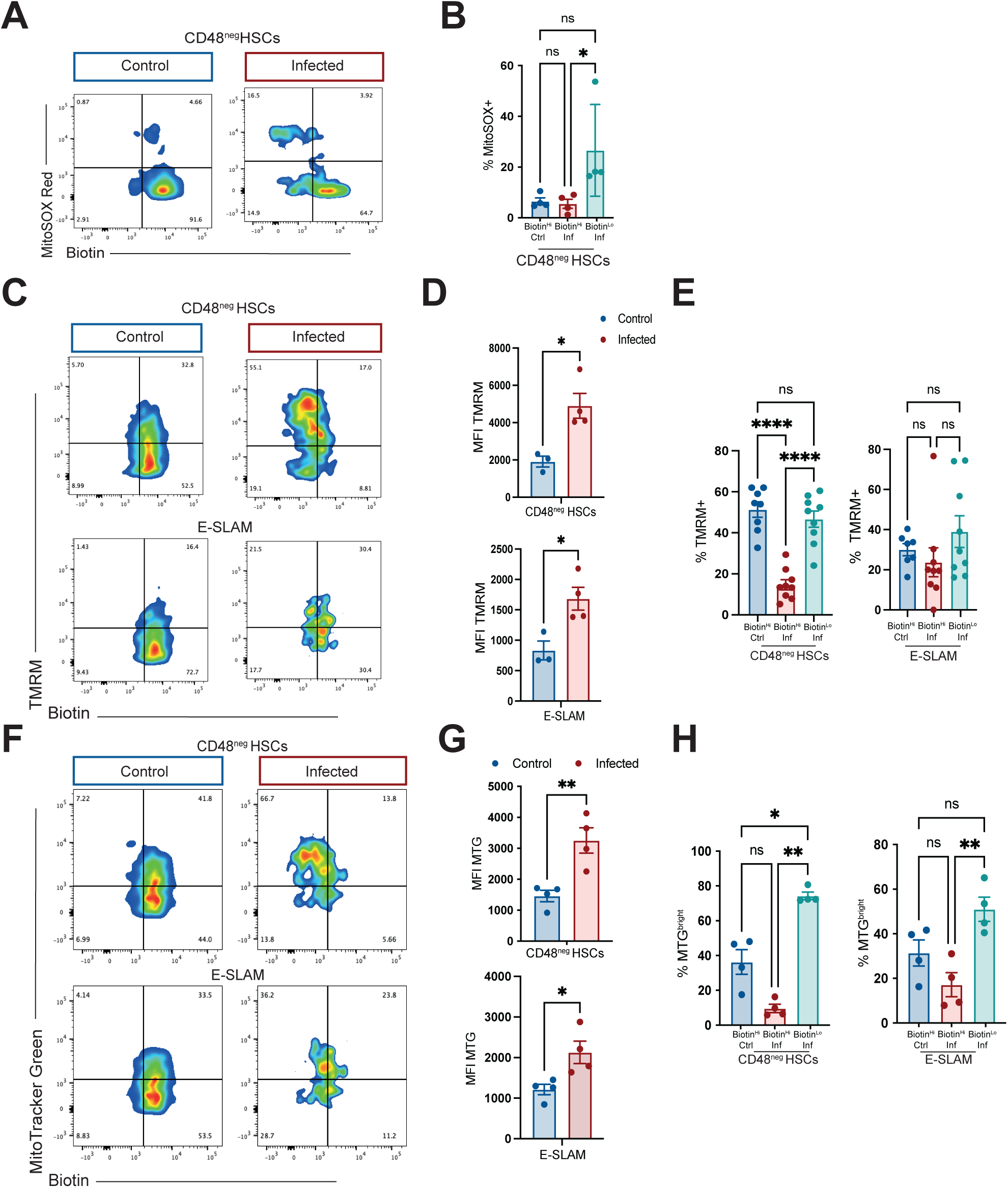
Biotin^Hi^ HSCs retain low mitochondrial activation. (A) Representative flow cytometry dot plot showing different mitochondrial ROS (MitoSox staining) in CD48^neg^ HSC Biotin subsets. (B) Percentage of MitoSOX labelled CD48^neg^ HSC Biotin subsets. In A and B n= 4 control and 4 infected animals from one of two representative infections. (C-E) TMRM labelling is used to assess mitochondrial mass in HSC subsets in control and infected mice. C shows representative dot plots of CD48^neg^ HSCs and E-SLAM HSCs from control and infected mice, D is quantification of TMRM MFI in the two populations (top and bottom, respectively) (n = 3 control and 4 infected mice from one representative infection), E is quantification of % TMRM+ cells in each Biotin subset in CD48^neg^ HSCs (left) and E-SLAM HSCs (right) n= 8 and 7 (CD48^neg^ and ESLAM analysis, respectively) control and 9 infected mice pooled from 2 independent infections. (F-H) Mito Tracker Green (MTG) is used to assess mitochondrial mass in HSC subsets. F shows representative dot plots of CD48^neg^ HSCs and E-SLAM HSCs from control and infected mice; G is quantification of MTG MFI in the two populations (top and bottom, respectively), H is quantification of % MTG bright cells in each Biotin subset in CD48^neg^ HSCs (left) and E-SLAM HSCs (right). n = 4 control and 4 infected mice from two independent infections. The data are presented as means ± s.e.m. ****p ≤ 0.0001;*** p ≤ 0.001** p ≤ 0.01; * p<0 05. The p values were determined by unpaired Student’s *t*-test and ordinary one-way ANOVA with Tukey correction.

Next, we measured mitochondrial activity in the HSC Biotin subsets using tetramethylrhodamine methyl ester (TMRM), reporting on MMP, and known to increase as a consequence of active OXPHOS (Figure 5C). TMRM uptake was clearly increased upon infection in unfractionated CD48^neg^ and E-SLAM HSCs populations, albeit significantly only in HSCs (Figure S4B and Figure 5C and Figure 5D), and in MPP 2 and 3 but not 4 populations (Figure S4C). Next, the proportion of cells showing MMP in infected Biotin subsets showed an increase in the Biotin^Lo^ HSCs, but not E-SLAM Biotin^Lo^, and a relatively high proportion of E-SLAM Biotin^Hi^ remained TMRM^neg^ (Figure 5E).

Lastly, we evaluated mitochondrial content using MitoTracker Green (hereafter MTG) which accumulates in the mitochondrial matrix (de Almeida et al., 2017; Filippi & Ghaffari, 2019; Vannini et al., 2016) (Figure 5F). When unfractionated HSC and E-SLAM populations per condition were analyzed we saw increased fluorescence in infected animals (Figure 5G and Figure S4D). It was not surprising to see higher MTG MFI in MPP2 and MPP3 (Figure S4E) as both MPPs activate during infection to proliferate, thus requiring energy. Because largely Biotin^Lo^ cells were actively cycling during infection, we noted an increase in the proportion of MTG^bright^ Biotin^Lo^ cells in both conditions (MTG^bright^ Biotin^Lo^) (Figure 5H). Taken together, our data indicate that infection-resistant HSCs maintain a low mitochondrial metabolic profile despite sensing infection and being more primed to proliferate than HSCs from healthy control mice.

## DISCUSSION

Haematopoietic stem cells’ heterogeneity has been described in multiple contexts. Lineage biased HSCs have been highlighted through single cell transplantation assays and barcoding and lineage tracing analyses (Carrelha et al., 2018; Dykstra et al., 2007a; Pietras et al., 2016; Rodriguez-Fraticelli et al., 2018, 2020; Schulte et al., 2015; Yamamoto et al., 2018). Heterogeneity in steady state proliferation dynamics was identified using label retention assays, which contributed to linking deep quiescence to maximum fitness (Wilson et al., 2015). HSC heterogeneity in the context of stress responses has been studied less. vWF+ platelet biased HSCs have been described to selectively respond to systemic platelet activation and depletion (Luis et al., 2023). Heterogeneity in the timing of reconstitution has been identified in transplantation assays of both murine and human HSCs (Kaufmann et al., 2021; Kucinski et al., 2024). While it is generally accepted that infection triggers the loss of quiescence and hinders long-term self-renewal and multipotency of HSCs, linked to transcriptional and metabolic changes (Ansó et al., 2017; Cabezas-Wallscheid et al., 2017; Mansell et al., 2021; Vannini et al., 2016), the existence and degree of heterogeneity in the HSCs’ response to infection-driven inflammation has been overlooked, leaving open the questions whether all HSCs may be equally damaged in these settings, and whether subpopulations may exist with varying degrees of activation, and through what mechanisms. To address this, we used *Plasmodium* blood-stage infection to gain a quantitative understanding of the heterogeneity within the HSC pool based on their proliferation kinetics, and characterised the resulting subsets in terms of their functionality, transcriptome and metabolic activation. While earlier studies showed that inflammation leads to loss of long-term label retaining HSCs (Bogeska et al., 2022; Essers et al., 2009), we use short-term dilution of NHS-ester-biotin (Nygren & Bryder, 2008; Luis TC et.al.,2023) to identify HSC subpopulations with differential proliferation dynamics as infection develops. Biotin^Lo^ HSCs dilute the label and very effectively uptake EdU, indicating that they likely proliferate more than once in the window from day 3 to day 5 post blood infection. Biotin^Hi^ HSCs retain the label, consistent with remaining quiescent; however, a higher proportion of Biotin^Hi^ HSCs from infected mice compared with Biotin^Hi^ HSCs uptakes EdU during the same time window, suggesting that while they have not divided, they are more likely to do so. A similar heterogeneity in proliferative dynamics was described in response to myeloablation (Bowling et al., 2020) and it is likely an evolutionarily conserved mechanism that allows preserving some HSCs from stress. The differences in cell cycle state between Biotin^Hi^ and Biotin^Lo^ HSCs from infected mice are echoed by their higher and lower fitness in serial transplantation settings. The higher serial reconstitution driven by Biotin^Hi^ HSCs from infected animals compared to Biotin^Hi^ HSCs from control animals is surprising, and suggests that either these HSCs remain highly functional despite being primed to proliferate, or that their strong reconstitution potential is driven by the fewer cells still in G0, which must be more functional than the equivalent, larger G0 control population.

Gene expression profiling of HSC Biotin subsets confirmed cell cycle skewing at the transcriptional level and surprisingly revealed that Biotin^Hi^ HSCs from infected animals are not shielded from inflammatory signals, but rather develop strong interferon response signatures, similarly to Biotin^Lo^ HSCs. Despite sensing and responding to inflammation, Biotin^Hi^ HSCs from infected mice unexpectedly maintained high stemness and repopulation scores. In this context, HSC function is therefore not an inverse correlate of IFN responses, but rather the result of uncoupling such responses from stemness attrition mechanisms. This is consistent with resistance to infection being not stochastic but rather deterministic, as proposed by another recent study (Munz et al., 2023). It is possible that HSCs are positioned at various depths of quiescence and respond to chronic infection accordingly, illustrating the complexity of HSC dynamics *in vivo*.

Single cell transcriptomic analysis of HSC subsets from control and infected mice provided some insights into the molecular mechanisms underpinning the high functionality of Biotin^Hi^ HSCs in infected animals. CD74 and an overall MHC II signature were highly expressed in this HSC subpopulation. While MHC II is expected to be upregulated in response to interferon, such upregulation was stronger in HSCs than in multipotent progenitors, highlighting a likely specific role in stemness maintenance. This is consistent with recent reports of high CD74 and MHC II expression in HSCs that remain highly functional and do not give rise to progeny following myeloablation (Bowling et al., 2020) driven by STAT1 (Li et al., 2022), which was too one of the top differentially expressed genes in Biotin^Hi^ HSCs in infected animals. Interestingly, expression of MHC II has recently emerged as an important immunosurveillance system that secures the integrity of the HSC pool (Hernández-Malmierca et al., 2022), and expression of the immune checkpoint-related CD112 was linked to latency of engraftment in human HSCs (Kaufmann et al.,2021), suggesting that complex interactions between HSCs and the immune system may underpin HSC heterogeneity in responding to multiple stressors.

Metabolic signatures suggesting higher levels of oxidative phosphorylation in Biotin^Lo^ HSCs in infected mice were consistent with multiple readouts of mitochondrial activity remaining low in Biotin^Hi^ HSCs, in agreement with reports that these changes are directly associated with HSC activation and differentiation (Hinge et al., 2020). Mitochondrial activity is intimately linked with quiescence and coupled with lysosomal activity (Liang et al., 2020). The low metabolic profile of Biotin^Hi^ HSCs is consistent with low lysosomal activity of human latent HSCs that resist regenerative stress in transplantation settings (Kaufmann et al., 2021; Laurenti et al., 2015). Whether intrinsic HSC heterogeneity pre-existing inflammation onset or cell-extrinsic signals allow Biotin^Hi^ HSCs in infected mice to uncouple interferon responses from metabolic activation remains an open question.

HSCs carrying mutations associated with clonal haematopoiesis of indeterminate potential (CHIP) are known to gain a selective advantage over wild type HSC clones in inflammatory environmental settings (Cai et al., 2018; Donato et al., 2023; Hormaechea-Agulla et al., 2021; Karpova et al., 2025; Zioni et al., 2023). None of the CHIP associated genes or downstream pathways were differentially expressed in Biotin^Hi^ HSCs from infected mice. We therefore propose that the mechanisms leading to the establishment of Biotin^Hi^ HSCs may become important targets for interventions aiming to retain healthy HSC functionality lifelong.

In summary, our data highlight heterogeneities in the HSC response to *Plasmodium* infection-induced inflammatory stress, and how this response is a multi-faceted process including inflammation sensing, metabolic activation, and cell division. These steps can be uncoupled from each other in HSCs that sense, yet withstand infection. These cells are therefore infection resistant, and interventions that would block metabolic activation may protect HSC function from infection-driven inflammatory stress.

## MATERIALS AND METHODS

### Mice

All animal work was carried out in accordance with the animal ethics committees at Imperial College London and Francis Crick Institute, and UK Home Office regulations (Animals Scientific Procedures Act, 1986) under license numbers 70/8403 and PP9504146. C57BL/6.J, mTmG (Muzumdar et al., 2007) and CD45.1 (C57BL/6.SJL) were bred at Francis Crick Institute. C57BL/6.N, Tuck-Ordinary (TO) and CD1 mice were obtained from ENVIGO UK.

Female mice > 6 weeks of age were used in all the experiments because they were more amenable to be co-housed within fewer cages. The mice were housed in Techniplast sealsafe plus (greenline) individually ventilated cages (IVCs) with appropriate enrichment and bedding. According to UK Home Office code of practice, the temperature, humidity and light cycles of where animals were housed were between 20 and 24 °C, 45-65% and 12h/12 h light cycle, with a 30-minute dawn and dusk period, respectively.

### Plasmodium berghei experimental model

The generation of infected blood stocks from mosquito bite infections was carried out as previously described (Haltalli et al., 2020; Vainieri et al., 2016). Briefly, TO mice were first treated with phenylhydrazine to induce reticulocytosis. Three days later, they were infected *i.p.* with fresh blood, containing 10^6^-10^8^ *P. berghei* ANKA wildtype clone 2.34 and exposed to naïve mosquitos to feed. At day 17 post blood feed, starved mosquitos carrying sporozoites in their salivary glands, were allowed to “bite back” on naïve animals. Passage 0 (P0) from these infected animals was harvested through cardiac puncture 5-6 days post mosquito feed and maintained in cryopreservation medium (containing 9 volumes Alsever’s solution with 1 volume of glycerol). To infect mice for the experiments, a donor mouse was injected with 5 x 10^7^ parasites *i.p.* at D-5. At D0 fresh P1 infected blood from this mouse was harvested through cardiac puncture, diluted in Alsever’s/glycerol medium (1 volume of blood and 2 volumes of Alsever’s solution) and injected 5 x 10^7^ parasites *i.p.* in the experimental cohort. Blood parasitaemia malaria was monitored from D3 post blood injection (pbi) until D5, when mice were sacrificed for analysis and before the onset of cerebral malaria symptoms.

### Parasitaemia measurement

To calculate the levels of parasitaemia in peripheral blood (PB) of infected animals, a blood smear taken from infected animals was methanol fixed for 20 seconds and stained with 1:10 Giemsa/water (Sigma Aldrich) for 20 minutes. Infected and uninfected Red Blood Cells (RBCs) are counted under a light microscope with 100x oil immersed objective.

### Drug treatments

To induce reticulocytosis for production of *P. berghei* infected mosquitoes, phenylhydrazine from a stock concentration of 6mg/ml (in PBS) was administered *i.p.* in mice at a dose of 10 μl/gm body weight three days prior to infection.

For N-acetylcysteine (NAC) treatment, NAC (Sigma-Aldlrich; 130 mg/kg) or PBS (vehicle control) were injected intraperitoneally daily, starting 5 days before infection and throughout until sacrifice and analysis.

EZ-link-NHS-Biotin-Sulfo-LCLCLC label (Luis et al., 2023; Nygren & Bryder, 2008) was injected intravenously (*i.v.)* at a concentration of 1 mg/6 g at d3 post blood injection (p.b.i) to study HSC proliferation kinetics as previously described. Label dilution was allowed for two days until the animal sacrifice on d5.

For thymidine analogue treatment, 5-ethynyl-2’-deoxyuridine (EdU) was administered chronically in drinking water from day 3 to day 5 pbi at a concentration of 0.3 mg/ml supplemented with 2% sucrose. In addition, on days 3 and 4 pbi animals received an intraperitoneal injection at a dose of 1 mg/ 200 μl, followed by a final intravenous injection of 1 mg /200 μl on day 5 pbi, 45 minutes before mouse sacrifice and analysis.

### Flow Cytometry

For haematopoietic cell analysis, bones were crushed in DPBS without calcium chloride (CaCl_2_) and magnesium chloride (MgCl_2_), with 5% fetal bovine serum. The cells were filtered through a 70 μM cell strainer, depleted of red blood cells and labelled with c-Kit (or CD117) magnetic MicroBeads (Miltenyi Biotech) enabling enrichment of c-Kit+ cells through the MACS Column technology (Miltenyi Biotech) using LS columns (Miltenyi Biotech). To phenotypically characterize HSCs, MPP subsets and E-SLAMs, the c-Kit enriched cells were immunostained with fluorophore-conjugated antibodies specific to mouse. All antibodies were from Biolegend and BD Biosciences unless specified, Lineage (CD4, CD8, CD5, Ter119, Gr-1 and B220/CD45), c-Kit (2B8), SCA-1 (D7), Flk2 (A2F10), CD150 (TC15-12F12.2), CD48 (HM 48-1), CD45 (30-F11), EPCR (RMEPCR1560(1560)), Streptavidin, MHC II. For more information on all the antibodies used see Supplementary Table 3.

EdU was detected and analyzed as previously described (Akinduro et al., 2018) using the Click-iT EdU kit (ThermoFisher).

For cell cycle analysis, cells were fixed and permeabilised using BD Cytofix/Cytoperm Fixation/Permeabilization solution kit according to manufacturer’s instructions and then stained with anti-Ki67 (B56) antibody overnight at 4°C with gentle agitation. The nuclear staining dye SYTOX Green was added to the cells 15 minutes before samples were analyzed.

For the analysis of Reactive oxygen species, CellROX FITC reagent was used according to the manufacturer’s instructions (Thermofisher Scientific) and as previously published (Haltalli et al., 2020).

For the analysis of mitochondrial stainings, cells were first enriched for c-Kit and stained for LKS-SLAM as above. Then to determine superoxide anions, mitochondrial mass and mitochondrial membrane potential, cells were washed and incubated at 37°C and 5 % CO_2_ for 30 minutes with MitoSox Deep Red (1μM Invitrogen), MitoTracker Green (25nM, Invitrogen) and TMRM (0.1μM, Invitrogen) respectively. Cells were washed and analyzed within 60 minutes of mitochondrial staining.

Calibrite beads (BD Biosciences) were used to determine absolute cell numbers in the population of interest as described previously (Haltalli et al., 2020)

All samples were acquired on BD Symphony A5 or BD LSR Fortessa flow Cytometers and all parameters were analyzed using FACS Diva and Flowjo (Tristar) (BD Bioscience) software.

### Transplantation assays

Tibias, femurs, ileac bones and sternum were harvested from mTmG or CD45.2 (C57BL/6) control and infected donor mice at day 5 pbi. The bones were crushed in DPBS (without CaCl_2_ and MgCl_2_ + 5% FBS + Penicillin/Streptomycin), filtered through a 70μM cell strainer and lysed to deplete RBCs. WBM from mTmG animals was labelled with a cocktail of biotinylated, FITC or BV510 or BUV395 conjugated lineage antibodies (CD5, CD11b, CD19, CD45R/B220, Ly6G/C, Gr-1, Ter119, CD4 and CD8) to isolate haematopoietic progenitor cells using the EasySep Mouse Hematopoietic Progenitor Cell Isolation Kit (Stem Cell Technologies Cat 19856A). Cells were purified through lineage depletion from mTmG donors or c-Kit enrichment from NHS-ester biotin injected donors. Resulting cells were immunostained for LKS SLAM markers and sorted on a FACS Aria III (BD Biosciences). 120 mTmG CD48^neg^ HSCs were transplanted into lethally irradiated (two doses of 5.5Gy, at least 2,5 hrs apart) CD45.1 recipients (per condition) alongside 300,000 support BM cells from CD45.1 donor. 10 HSCs per Biotin type (high or low) per condition were transplanted into lethally irradiated CD45.1 recipients together with 250,000 support WBM from CD45.1 donor. Transplanted mice were monitored every 4 weeks up to week 24 by flow cytometry analysis of peripheral blood and, at the last time point, of whole bone marrow. For secondary transplants, 1 x 10^6^ WBM cells from CD45.1 primary recipients that received either 120 mTmG HSCs or 10 E-SLAM Biotin high/low from infected or control donors were injected into lethally irradiated CD45.1 secondary recipients. Baytril antibiotic was administered to recipient mice for 5 weeks post-transplant in both primary and secondary transplants.

### Single Cell RNA sequencing and analysis

Single cell RNA-Seq was performed as previously described (Picelli et al., 2013; Wilson et al., 2015). Briefly, single c-Kit enriched LKS SLAM HSCs purified from control and infected animals were prepared as described above and index sorted using a FACS Aria III (BD Biosciences) into individual wells of 96-well plates containing 2.3 μl of lysis buffer (SUPERase-In RNAse Inhibitor-Ambion AM2694, 10% Triton X-100-Sigma Aldrich 93443, RNAse free H_2_O-Qiagen). Plates were centrifuged and stored at –80°C. After thawing, 2 μl of the annealing solution (0.1 μl of ERCC RNA Spike-In solution (1:300,000 dilution), 0.02 μl of the Oligo-dT (100 μM stock concentration), 1 μl of dNTP (10 mM stock concentration) and 0.88 μl of dH_2_O) was added. The plate was incubated at 72°C for 3 minutes, immediately cooled on ice and reverse transcription was performed by adding 5.7 μl of annealing mix containing 0.1 μl of Maxima H Minus (200 U/ μl), 0.25 μl of RNAse Inhibitor (20 U/ μl), 2 μl of 5 x Maxima RT Buffer, 0.2 μl of TSO (100 μm), 1.875 μl of PEG 8000 (40%v/v) and 1.275 μl of dH_2_O then incubated at 42°C for 90 mins, followed by 70°C for 15 mins. 40 µl of PCR mix was added to each well (1 μl of Terra PCR direct Polymerase (1.25 U/ μl), 25 μl of 2x Terra PCR Direct Buffer, 1 μl of IS PCR primer (10 μM stock concentration) and 13 μl of dH_2_O) and the cDNA was amplified by PCR, using the following PCR conditions: 98°C for 3 min, followed by 98°C for 15 s, 65°C for 30 s, 68°C for 4 min for 19 cycles, 72°C for 10 min. Then, amplified cDNA was purified using AMPure XP Beads and quantified using the Scientific Quant-iT PicoGreen dsDNA Assay kit. For library preparation, 100-150 pg/ μl of cDNA was used along with 2.5 μl of Tagment DNA buffer and 1.25 μl of Amplicon Tagment Mix (Illumina Nextera XT Kit). Samples were incubated at 55°C for 10 min and 1.25 μl of NT buffer was added immediately after samples reached 10°C to neutralise the reaction. Index Primer 1 (N701-N712) and Primer 2 (S501-S508) were added to tagmented cDNA and then amplified at 72°C for 3 min, 95°C for 30 secs, 12 cycles of (95°C for 10 s, 55°C for 30 s, 72°C for 60 s), 72°C for 5 min. Libraries were sequenced using the Illumina Hiseq4000.

### Data processing and quality control

In total, there were 288 cells from three different plates. First, the data were mapped to the mm10 reference genome with 92 spike-ins developed by the External RNA Control Consortium (ERCC) using STAR. The raw expression matrix was then generated using featureCounts.

Quality control and normalisation were performed using the in-house software smqpp (https://github.com/SharonWang/smqpp), which was constructed to filter out some low-quality cells in Smart-seq2 data. Briefly, based on several features after mapping, including the number of reads mapped to mitochondrial genes (nMito), the number of reads mapped to nuclear genes (nNuclear), the number of reads mapped to ERCC spike-ins (nERCC), the number of high-coverage genes (count per million greater than 10) (nHCGenes), the total number of mapped reads (nMapped), the total number of reads (nTotal), a cell failing one of these followed criteria was removed (nMapped>0, nNuclear>10^5.2^ (nMito+nNuclear):nTotal>0.2, nHCGenes>4000, nMito:nNuclear<0.2, nERCC:nMapped<0.4). 113 cells were removed in this process. In addition, 2 cells without biotin information were removed from the subsequent analysis. Thus, a total 165 cells were retained. Furthermore, genes that were not expressed in any cell were deleted using the scanpy.pp.filter_genes function. As a result, a total of 28,353 genes were filtered. Finally, the expression matrix containing the raw count values for all 165 selected cells with 27,061 genes was normalized to a total count of 10,000 in cell-wise and log-transformed using the smqpp function normalise_data.

#### Identification of variable genes

Highly variable genes (HVGs) were detected using the tech_var function in smqpp, which follows Brennecke’s approach. Default parameters were used here and ERCC was not considered (meanForFit=10, useERCC=False).

#### Clustering and mapping

Uniform Manifold Approximation and Projection (UMAP) was used for visualization using the umap function in Scanpy. The normalized data containing all selected HVGs was first scaled using the scanpy.pp.scale function, and then the top 50 principal components were calculated using the scanpy.tl.pca function. Considering the variance ratio explained by each principal component and not involving too much noise, a neighborhood graph was computed using the first 15 components and the number of nearest neighbor cells was set to 11.

#### Cell-cycle analysis

Normalized and log-transformed data was scaled before calculating cell cycle scores. The three phases of the cell cycle (G0/G1, S, and G2M) were estimated using the scanpy.tl.score_genes_cell_cycle function using 97 genes associated with cell cycle signature from (Kowalczyk et al., 2015).

#### hSCscore, p53, aging and repopulation scores

All four measurement scores are calculated based on the expression values of a subset of genes defining a specific signature. The HSC Score is calculated based on the expression values of the 103 molecular overlapping population (MolO) signature genes from our previous study(Hamey & Göttgens, 2019; Wilson et al., 2015). The hematopoietic p53 Score is calculated based on the expression values of 16 *Trp53* target genes (Wang et al., 2023). The Aging signature score was calculated based on the expression values of the best 20 age-associated genes from 220 aging signature genes (Arthur Flohr Svendsen, 2021). The Repopulation signature score was calculated based on the expression values of 23 relevant Repopulation signature genes (Che et al., 2022). In addition to calculating the HSC Score using the hscScore model (Hamey & Göttgens, 2019), the p53Score, Aging score and repopulation signature score were all generated using the score_genes function in Scanpy.

#### Differential gene expression analysis

Differential expression analysis between two categories was performed using rank_genes_groups function in scanpy. Differentially expressed genes (DEGs) were detected according to the thresholds set by adjusted p-value and log fold changes (adjusted p-value<0.05; absolute value of logfoldchange>1.5). Volcano plots were utilized to demonstrate the DEGs. Additionally, significant DEGs were used for Gene Ontology enrichment analysis by Enrichr. All plotting was performed using ggplot2 package in R.

#### MHC class II core enrichment score

All differentially expressed genes in the Infected Biotin^Hi^ and Control Biotin^Hi^ cells were first ranked. The ranked values were derived from the multiplication of the log fold change with the negative log 10 of the adjusted p-value. The ranked genes with the ranked values were then used as inputs to the GSEA software to perform Gene Set Enrichment Analysis using the pre-rank mode. The top 6 core enrichment genes for MHC class II signatures reported in GSEA were then selected and gene scores were calculated using the scanpy score_genes function.

#### Metabolic flux analysis of Glycolysis and TCA cycle reactions

The method constructed in our previous study was utilized directly here (Isobe et al., 2023). Briefly, normalized expression counts were used as input for the python package scFEA (Alghamdi et al., 2021). Metabolic flux values for the 168 core metabolic reactions implemented in scFEA were inferred using the default parameters. The inferred metabolic flux values were compared between the Biotin^Hi^ and Biotin^Lo^ cells by logistic regression and likelihood ratio test. The size of differences between two groups was evaluated by Cohen’s D standardized mean differences. The metabolic reactions with |Cohen’s D| > 0.15 and Benjamini-Hochberg (BH)-adjusted p values < 0.05 were considered significant.

### Statistics and reproducibility

Raw data were visualized and processed using Microsoft Excel and GraphPad Prism (GraphPad Software). Group means were compared using unpaired, Student’s *t*-tests. For multiple comparisons, unpaired, one-or two-way analysis of variance (ANOVA) with post-hoc Bonferroni or Tukey corrections were used. For all data, differences were considered significant when p<0.05.*p<0.05;**p<0.01;***p<0.001;****p<0.0001. Specific statistical parameters (e.g., number of animals used) can be found in the figure legends. All specific statistical details can be found in the figure captions. For all experiments, *n* corresponds to the number of independent measurements providing values for statistical analysis. For biological replicates, at least 2 independent infections were performed for all *in vivo* experiments, with similar results obtained. Animals were randomly allocated to control or experimental conditions.

## ACKNOWLEDGMENTS

This work was funded by Wellcome, Cancer research UK and Royal Society (Investigator award 212304/Z/18/Z, Programme Foundation Award C36195/A26770, and International Exchange Grant IEC\R1\180061, respectively, to CLC). SGA was funded by a Cancer Research UK PhD studentship (C36195/A27830). FBruno was funded by an ICL DoLS PhD studentship. FBirch was funded by Fondation Alcea (personal fellowship, and project grant to CLC). Work in the Göttgens laboratory is supported by Wellcome (206328/Z/17/Z), Blood Cancer UK (18002), Cancer Research UK. Part of this work was conducted at the Cambridge Stem Cell Institute, which is supported by the Wellcome Trust (203151/Z/16/Z, 203151/A/16/Z) and the UKRI Medical Research Council (MC_PC_17230). TCL was supported by a Wellcome Trust Sir Henry Dale Fellowship (210424/Z/18/Z) and a Kay Kendall Leukaemia Fund Project Grant (KKL1379). AMB thanks the MRC (MR/N00227×/1and MR/W025701/1), Sir Isaac Newton Trust, Alborada Fund, Wellcome Trust ISSF and University of Cambridge JRG Scheme, GHIT, Rosetrees Trust (G109130) and the Royal Society (RGS/R1/201,293) (IEC/R3/19,302[HR1]). Part of this work was conducted at the Sir Francis Crick Institute, which is supported by Wellcome, Cancer Research UK and the UKRI Medical Research Council. We thank K. Sala for passaging parasites, preparing mosquitos and advice with infections and all of the Lo Celso group members for constructive discussions. For the purpose of open access, the author has applied a CC BY public copyright licence to any Author Accepted Manuscript version arising from this submission.

We thank Nick Van Gastel and Richard Burt for critical discussions and feedback, Ilaria Malanchi and Dominique Bonnet for their overall support, and Ken Duffy for discussion of statistical analyses. We thank Imperial College Flow Cytometry and Central Biomedical Services facilities, and the Sir Francis Crick Flow Cytometry facility and Biomedical Research Facility for their technical support.

## SUPPLEMENTARY FIGURE LEGENDS

**Supplementary Figure 1.**
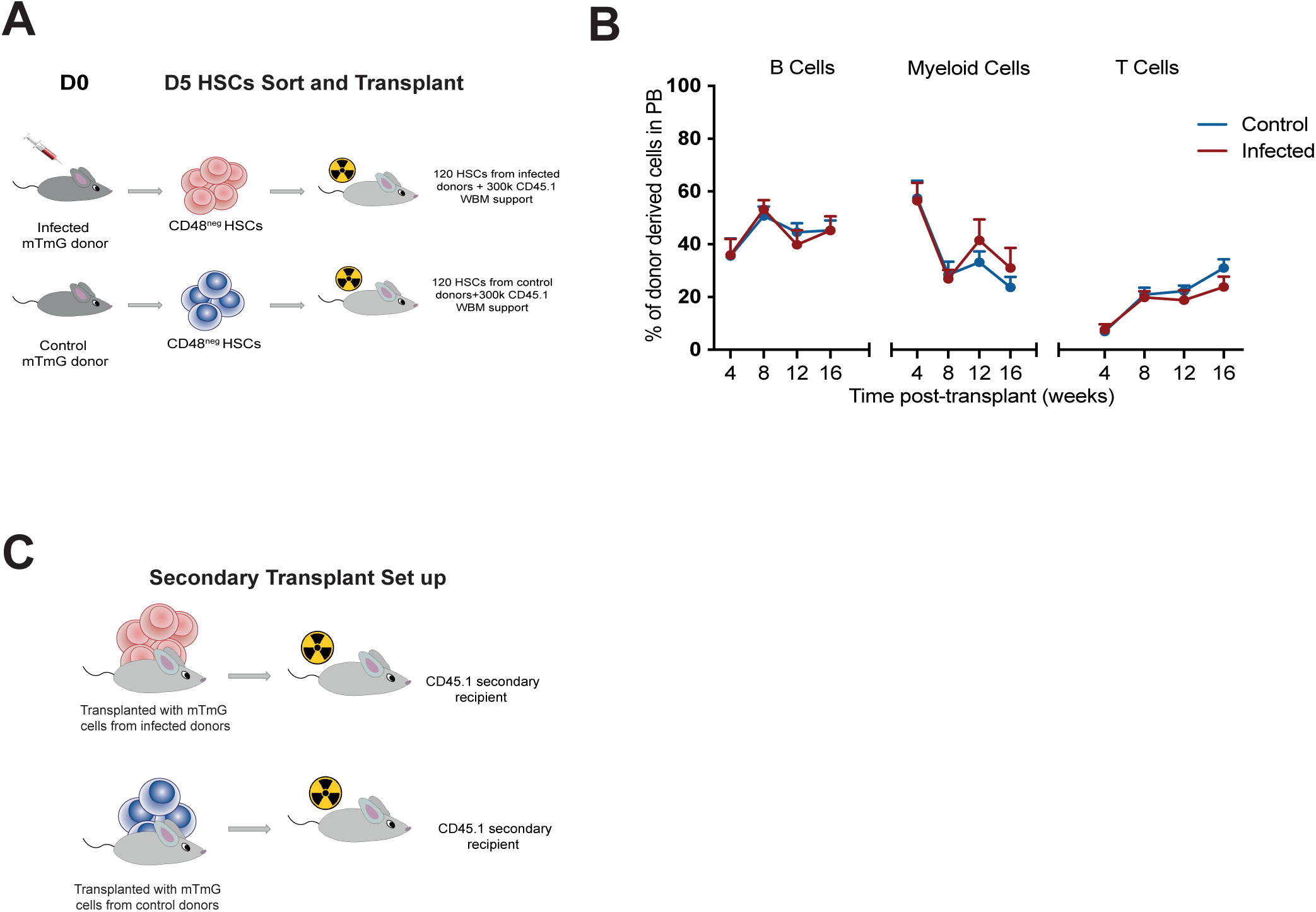
(A) Experimental set up. 120 CD48^neg^ HSCs were sorted from control and infected mTmG+ donors and were injected into lethally irradiated CD45.1 recipients together with 300,000 CD45.1 WBM MNCs support. (B) Multilineage reconstitution of mTmG^+^ HSCs expressed as a percentage of lineage output (T, B, myeloid cells) in PB assessed over 16 weeks. n= 9 recipient mice per group, pooled from two independent experiments. Data are presented as means ± s.e.m. The p values were determined by two-way ANOVA with post-hoc Bonferroni correction. No statistically significant differences were identified.(C) Experimental set up of secondary transplants where 1M WBM cells from pooled primary recipients were injected into lethally irradiated CD45.1 recipients. Engraftment was monitored over 16 weeks as the percentage of mTmG+ cells in PB. In g and h n= 4 and 5 recipients of WBM from primary recipients in control and infected groups, pooled from 2 independent experiments. In B statistically significant differences were determined using two way ANOVA with post-Tukey correction (**** represents p≤ 0.0001; *** represents p≤0.001; ** represents p≤0.01; * represents p≤0.05). Error bars represent mean ±s.e.m.

**Supplementary Figure 2.**
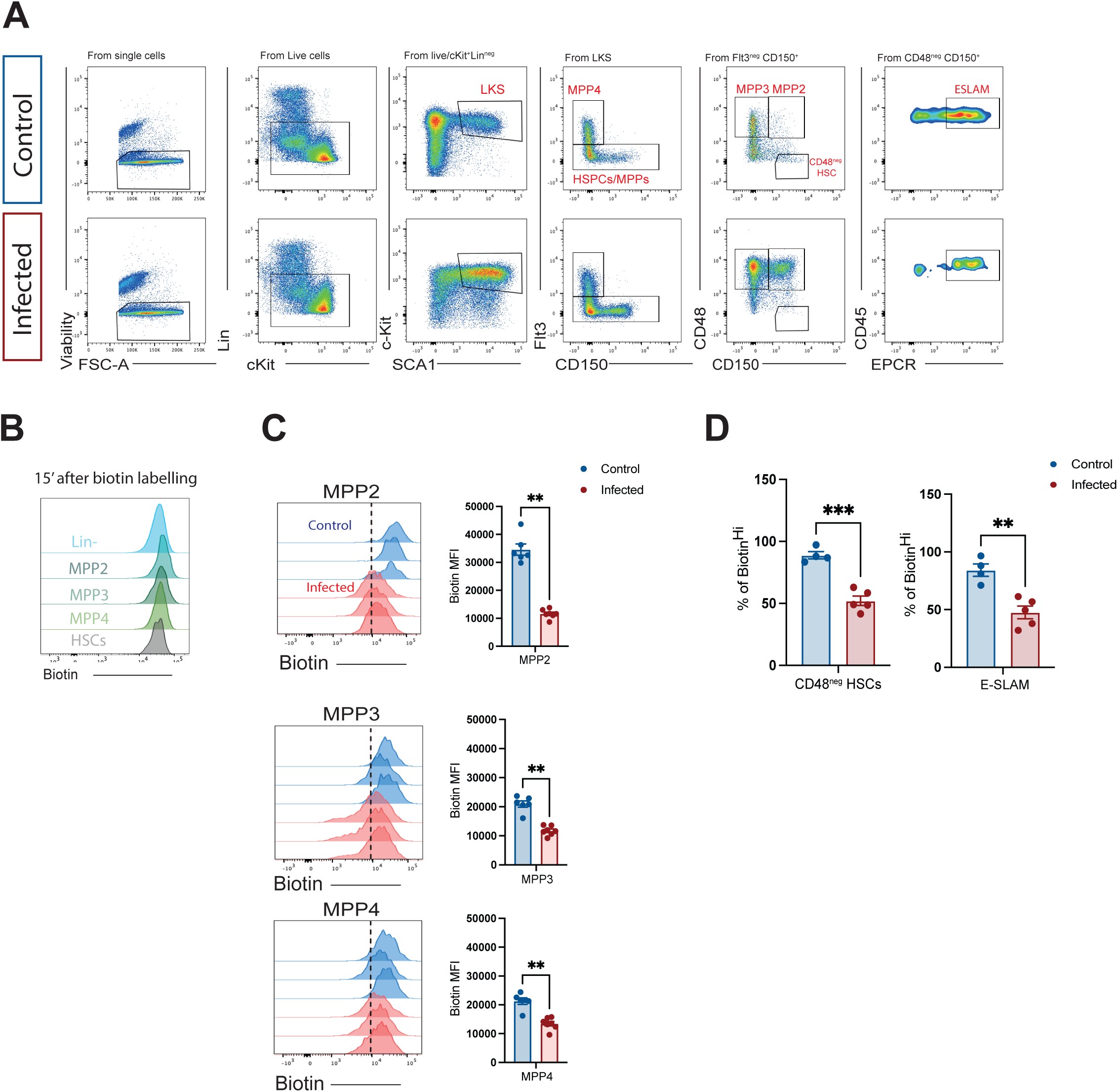
(A) Gating strategy showing CD48^neg^HSCs, E-SLAM HSC subset and MPP populations. (B) Representative histograms showing N-hydroxysulfosuccinimide (NHS)-ester-LC-LC-biotin (NHS-ester-biotin) labelling in Lineage^neg^ cells, MPP populations and HSCs in control animals sacrificed 15 minutes after *i.v.* injection. n= 5 animals pooled from two independent experiments. (C) NHS-ester Biotin dilution in the indicated MPP populations from control and infected mice. Representative histograms on the left (Dotted line represents the threshold between Biotin^Hi^ and Biotin^Lo^ cells), and biotin label MFI quantification on the right. n=6 animal per group, pooled from 2 independent infections. **: p,0.001 calculated with unpaired t-test. Data are presented as means ± s.e.m. ** p ≤ 0.01. The p values were determined by unpaired Student’s *t*-test.

**Supplementary Figure 3.**
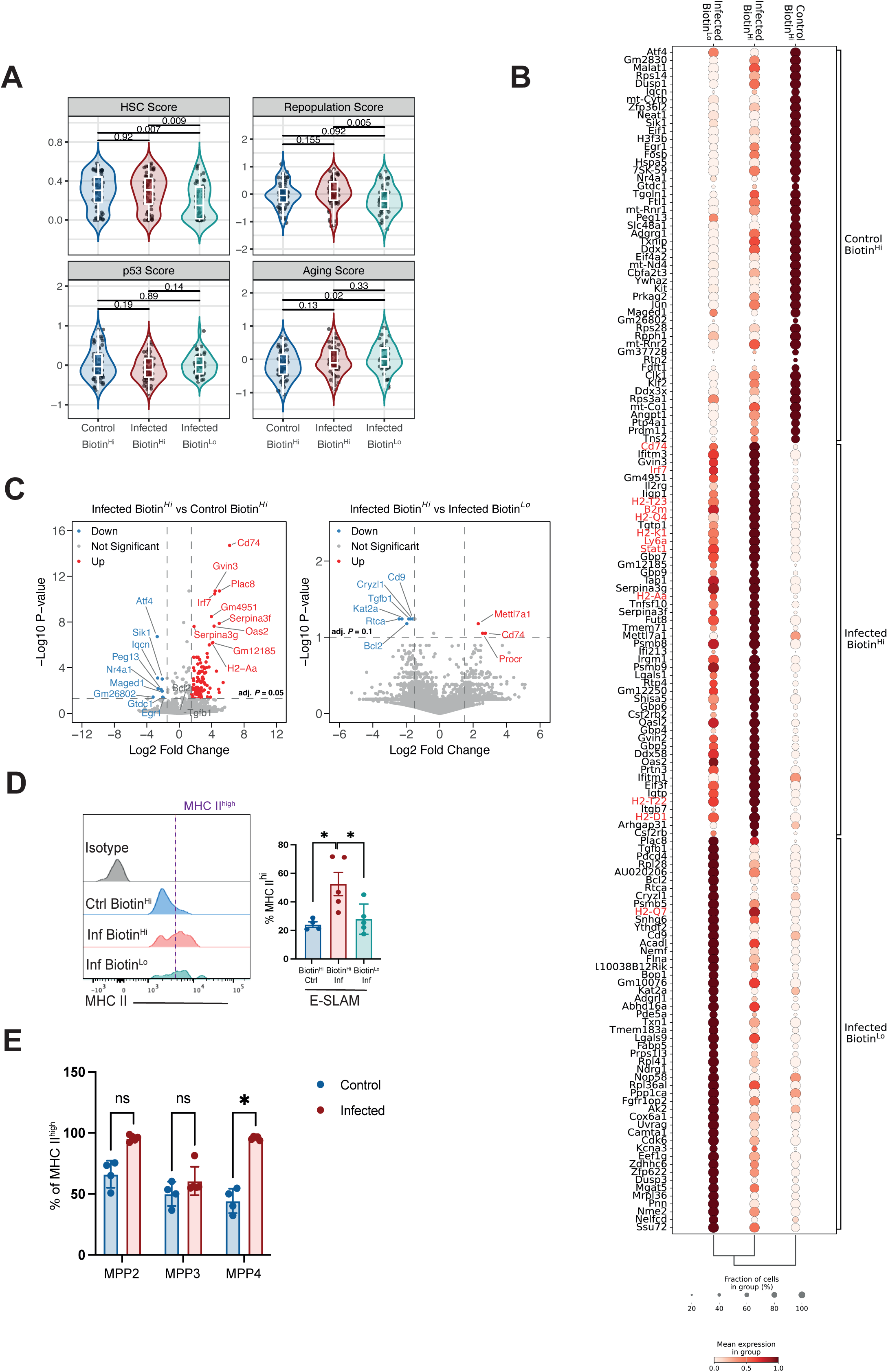
(A) HSC, repopulation, p53 and aging scores values for Biotin^Hi^ control, Biotin^Hi^ infected and Biotin^Lo^ infected cells. n: see Figure 4A. The p values were determined using Wilcoxon test. (B) Dotplot highlighting the top 50 genes differentially expressed in each category of cells. Genes in red are found in the GO categories ‘Cellular response to type II interferon’ and ‘Response to type II interferon’ shown in Fig. 4F. (C) Volcano plot highlighting Differentially Expressed Genes (DEGs) between Biotin^Hi^infected and Biotin^Hi^ control (top) and between Biotin^Hi^ and Biotin^Lo^ infected (bottom) cells. Highlighted in colour are genes that are over 1.5 fold up/downregulated and with adjusted p value <0.05 (top) and <0.1 (bottom). (D-E) Representative Flow cytometry analysis of MHC II expression in E-SLAM HSC (D) and MPP (E) populations. n= 8 control and 9 infected mice from two independent infections. The data are presented as means ± s.e.m. The p values were determined by unpaired Student’s *t*-test or ordinary one-way ANOVA with Tukey correction.* p<0 05.

**Supplementary Figure 4.**
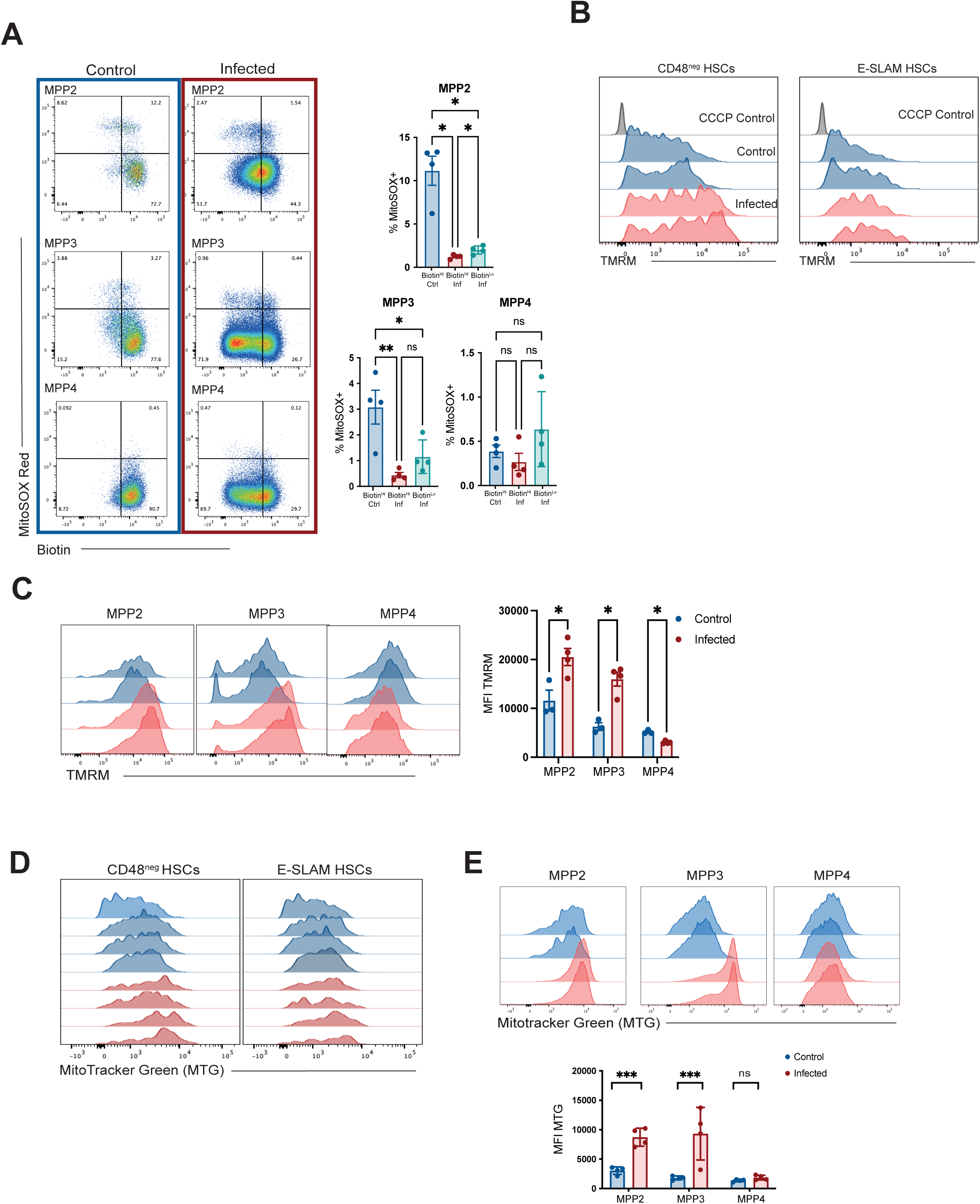
(A) MitoSOX staining of MPP populations from control and infected mice. Left: representative dot plots of the populations indicated, showing MitoSOX vs Biotin label dilution. Right: % MitoSOX positive cells in the biotin subsets of the MPP populations indicated. n = 4 control and 4 infected mice from 1 of 3 independent infections. (B) Representative histograms showing TMRM staining in CD48^neg^ HSCs and E-SLAM HSCs from control and infected mice. n = 3 control and 4 infected mice from one representative infection. (C) TMRM staining of MPP 2, 3 and 4 populations. Left: representative histograms. Right: quantification of MFI values. n = 3 control and 4 infected mice from one representative infection. (D) Representative histograms of Mitotracker Green (MTG) staining in CD48^neg^ and E-SLAM HSCs. n = 4 control and 4 infected mice from two independent infections. (E) Representative histograms of Mitotracker Green (MTG) staining in MPP populations (top) and MFI quantification (bottom). n= 4 control and 4 infected mice from two independent infections. Data are presented as means ± s.e.m. ****p ≤ 0.0001;*** p ≤ 0.001** p ≤ 0.01; * p<0 05. The p values were determined by unpaired Student’s t-test or ordinary one-way ANOVA with Tukey correction.

---

**Supplementary Table 1.**
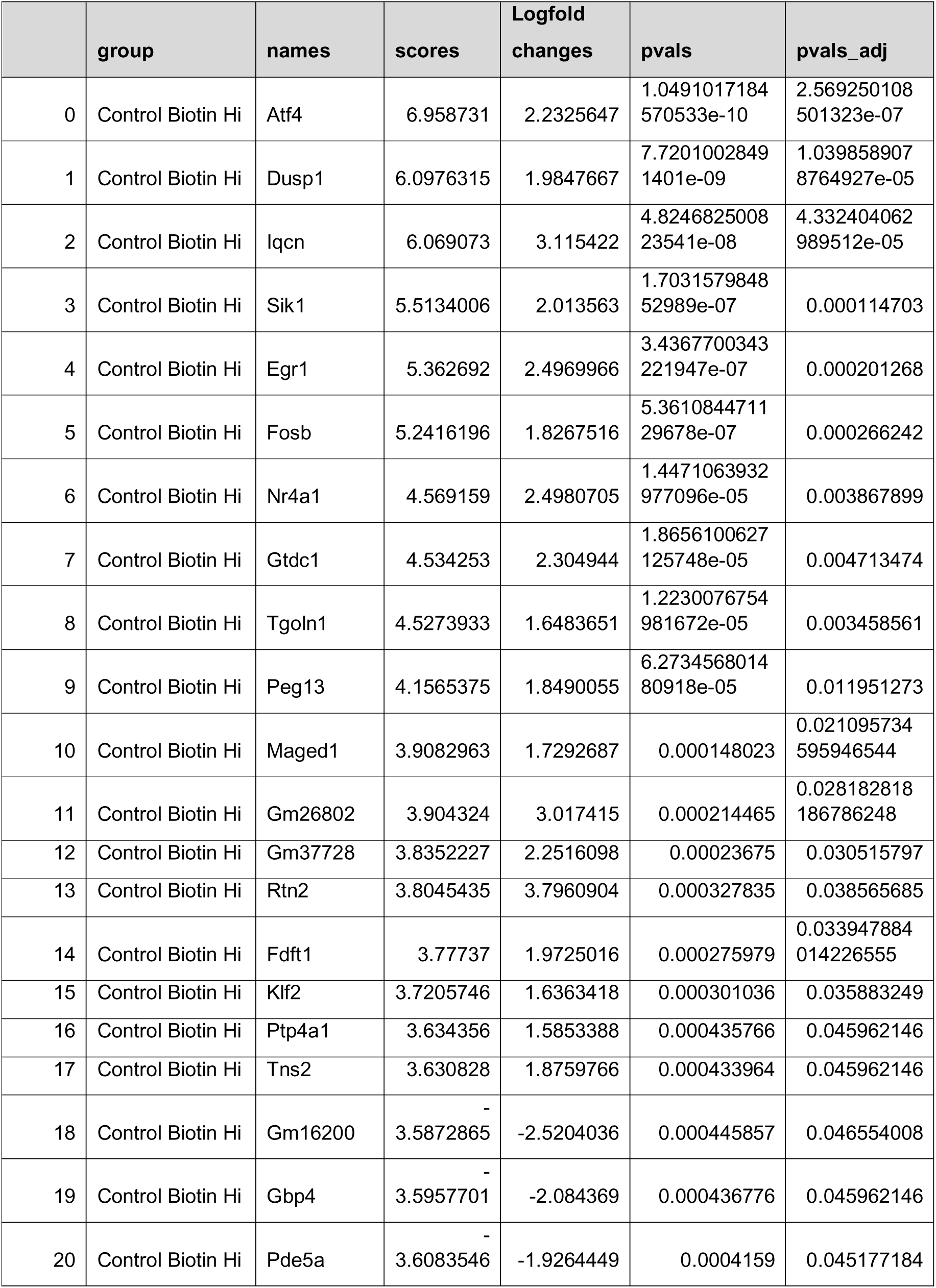
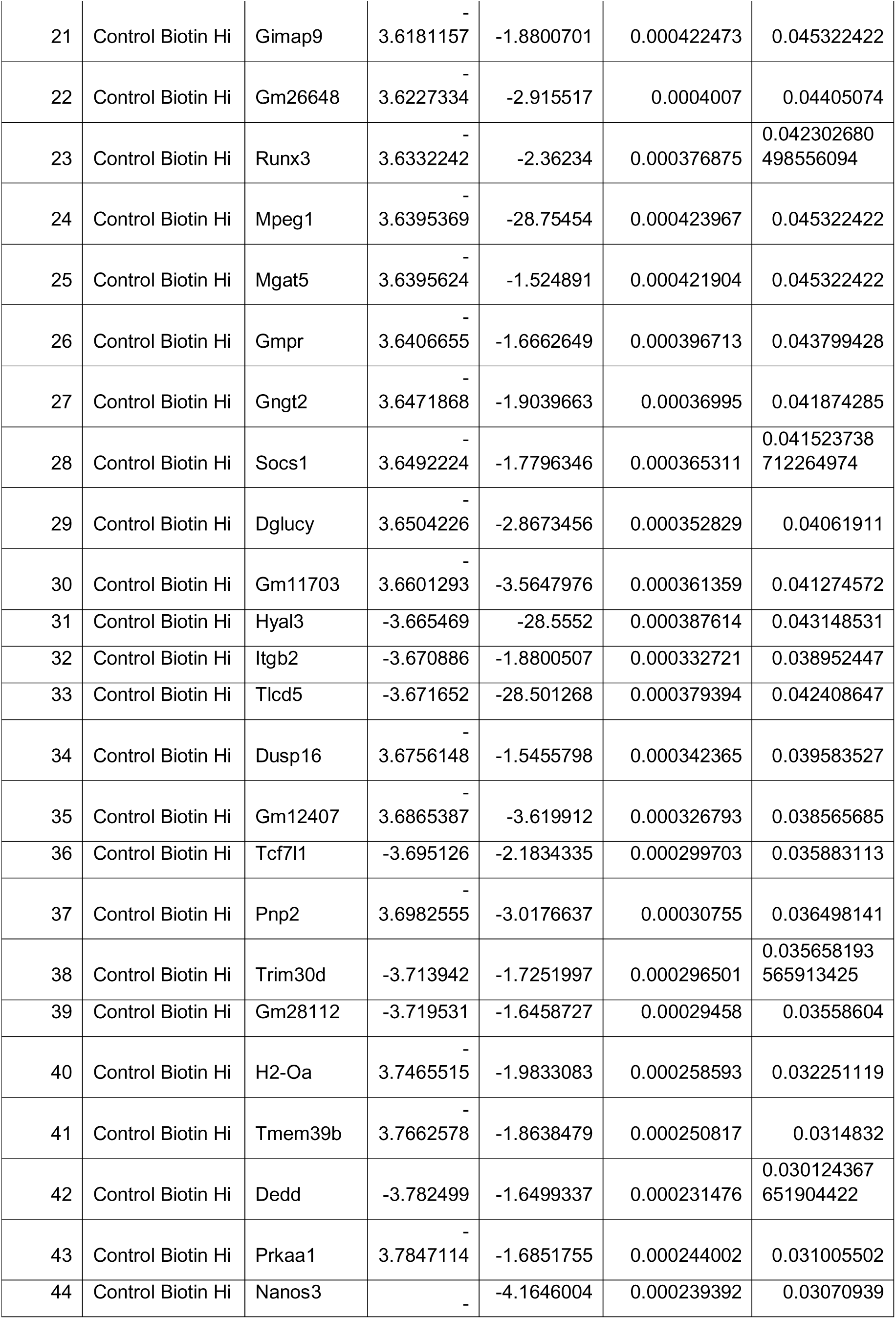
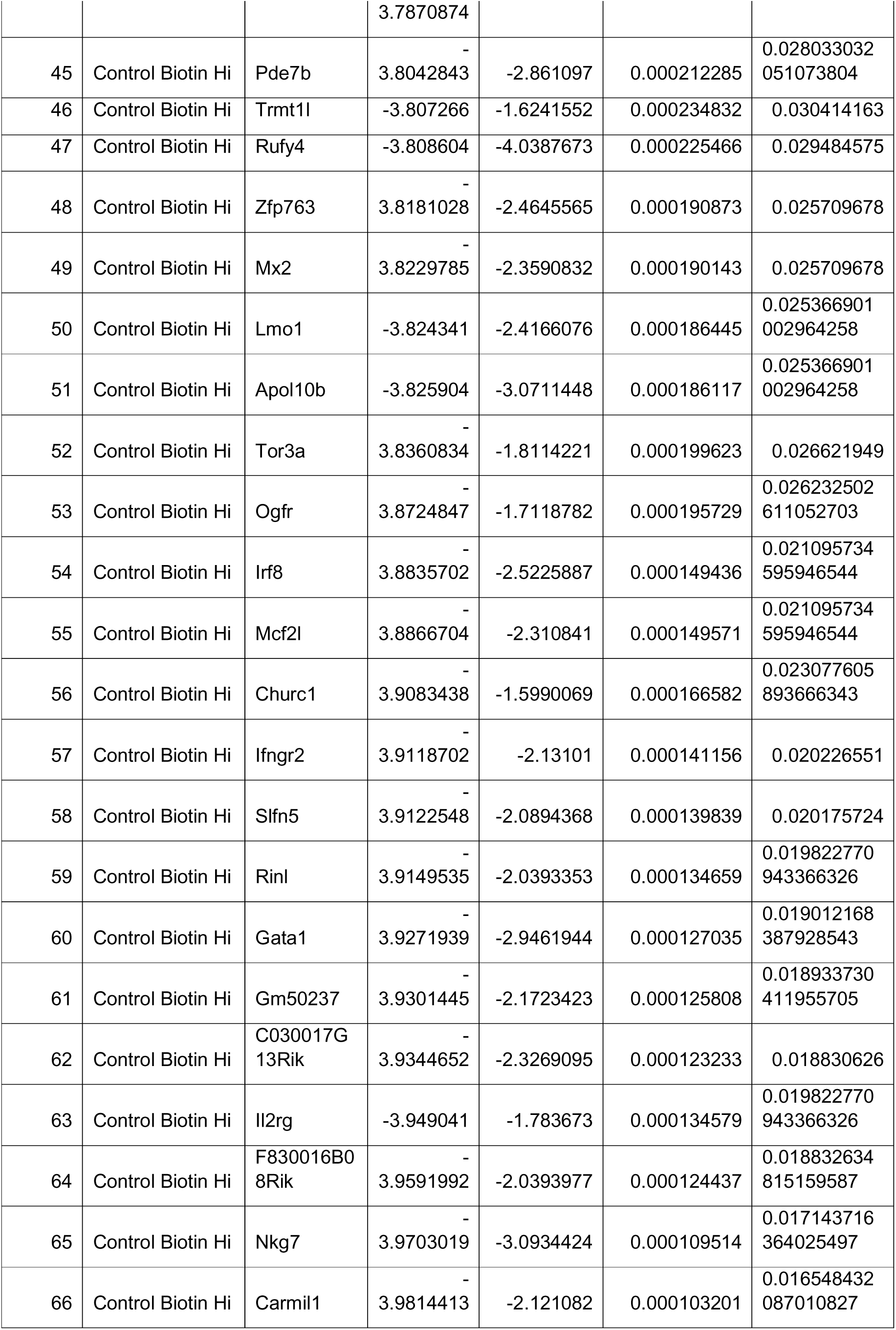
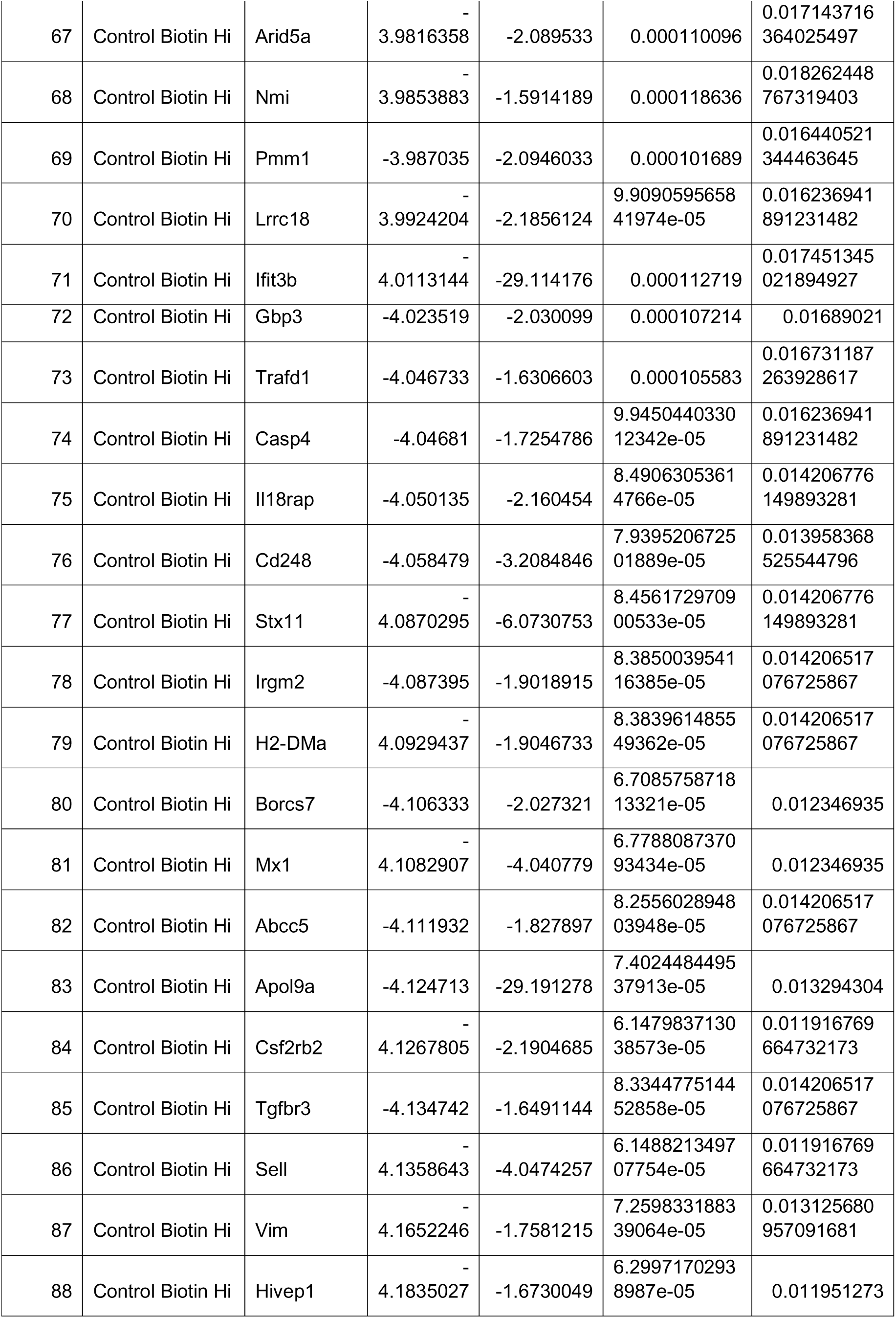
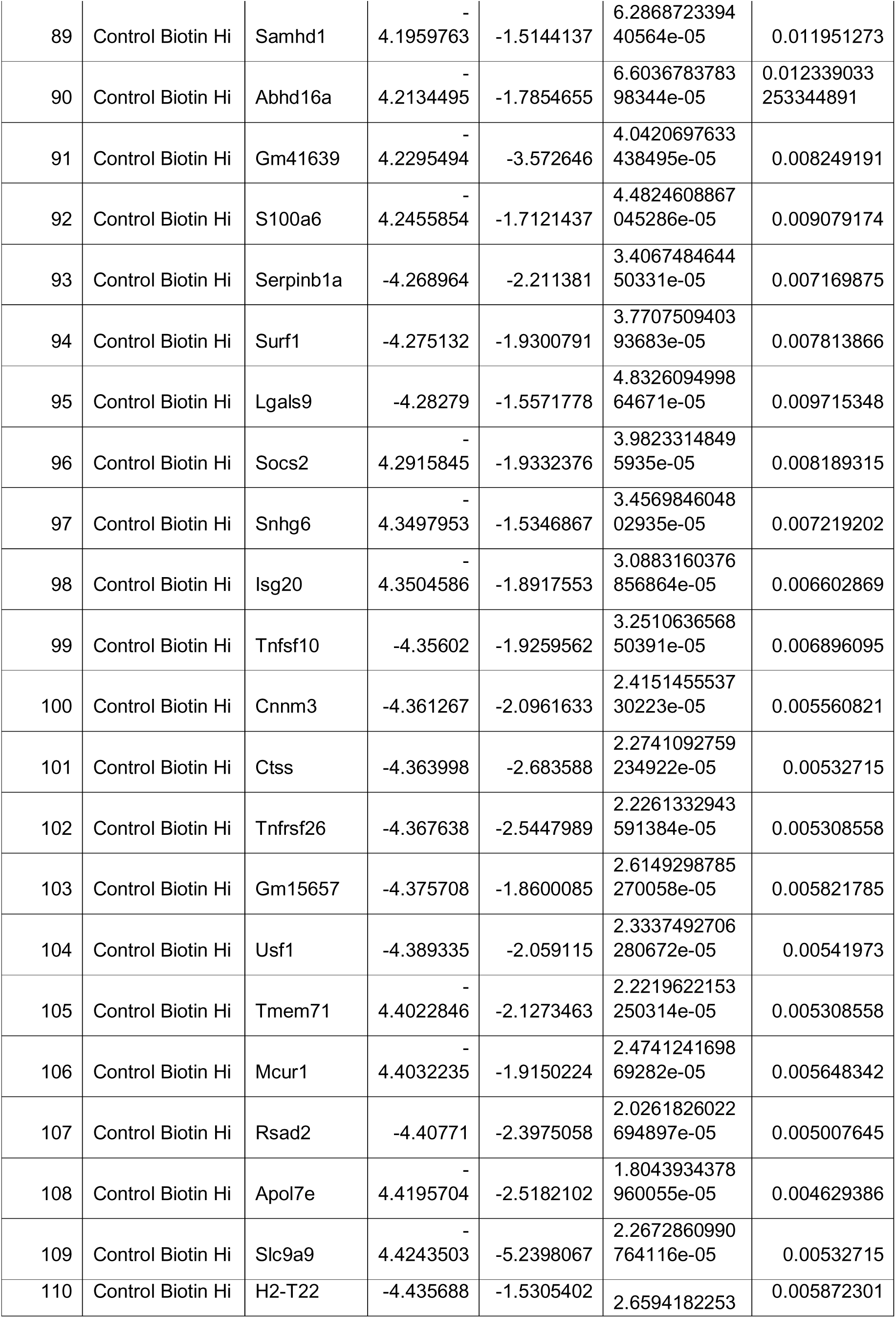
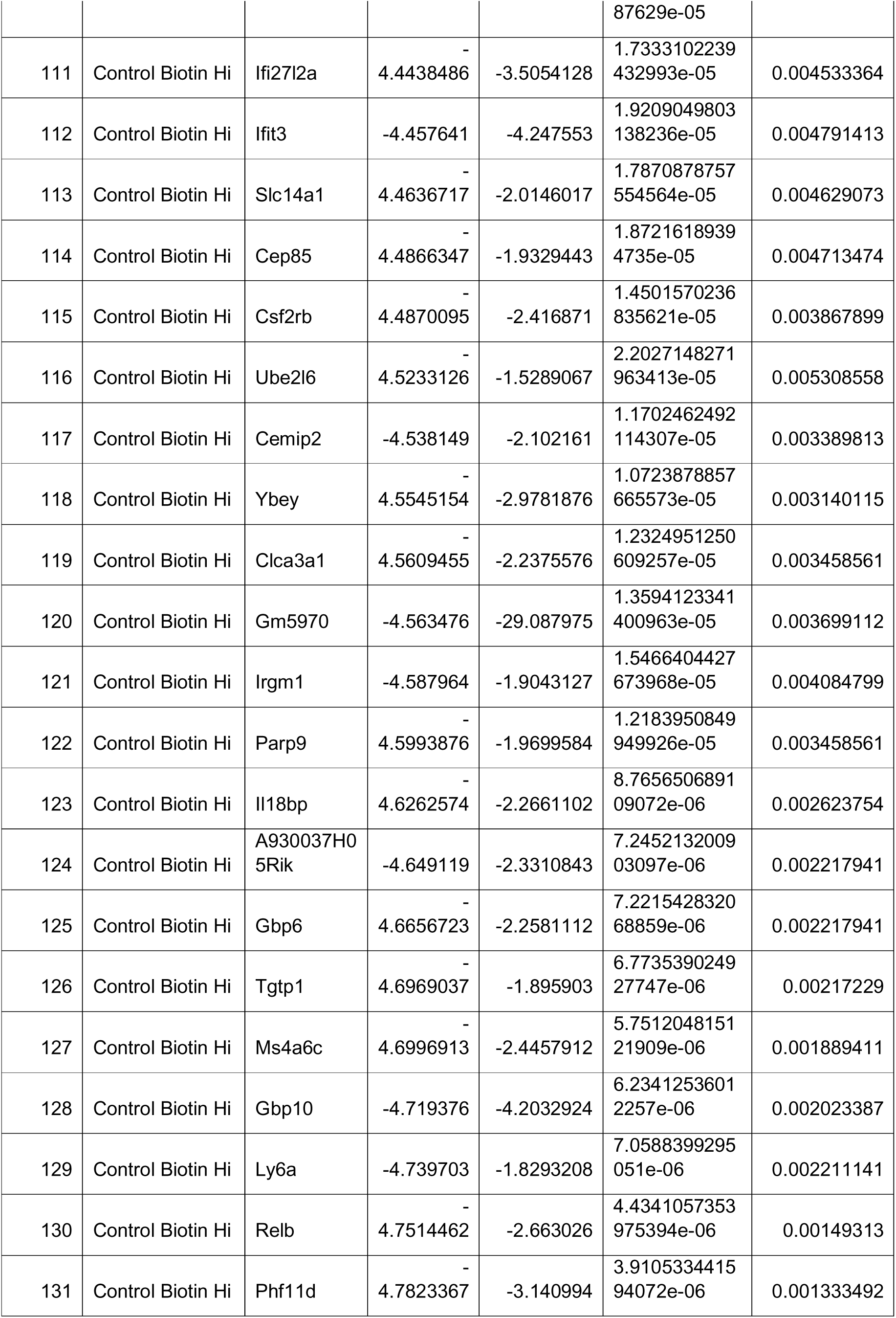
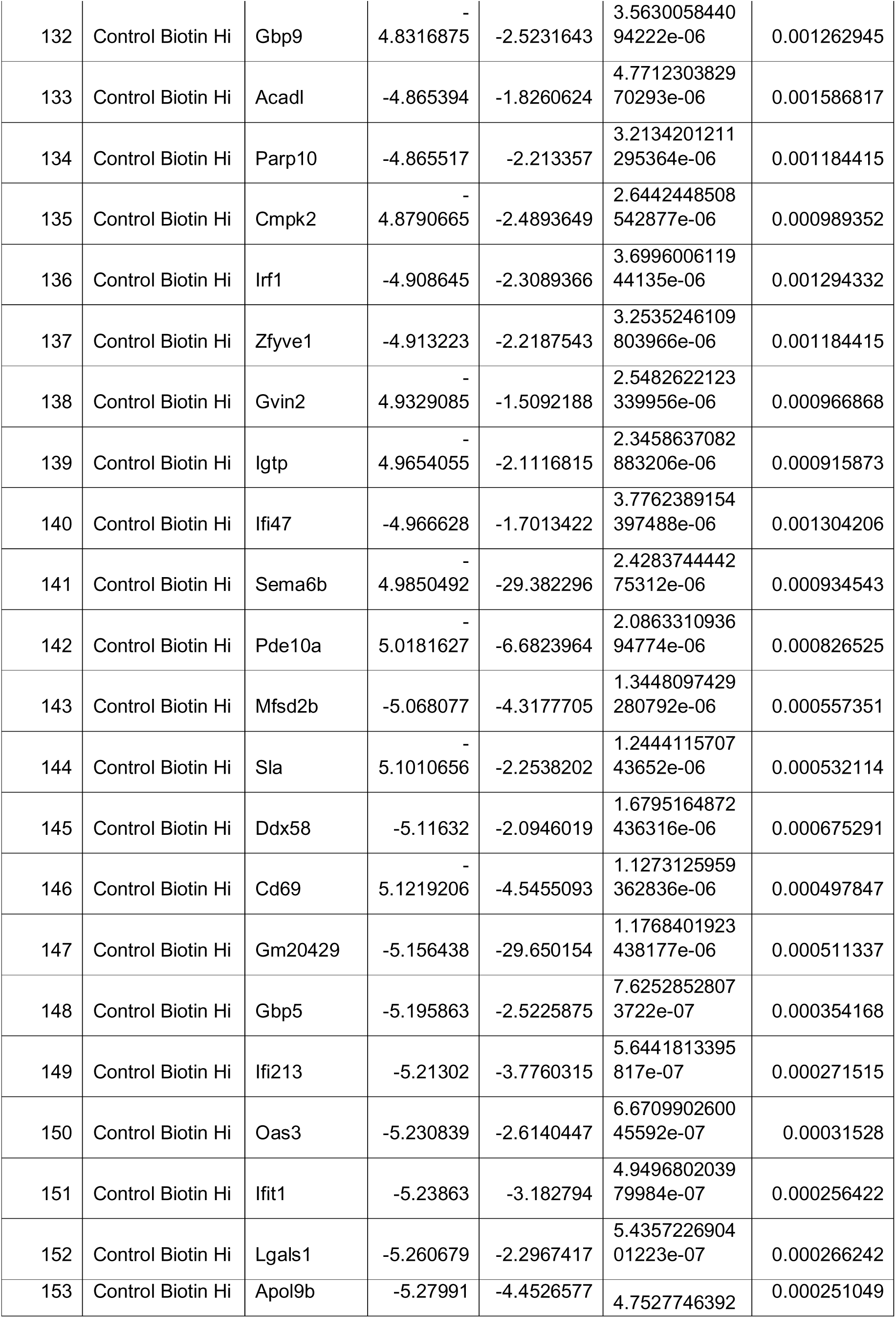
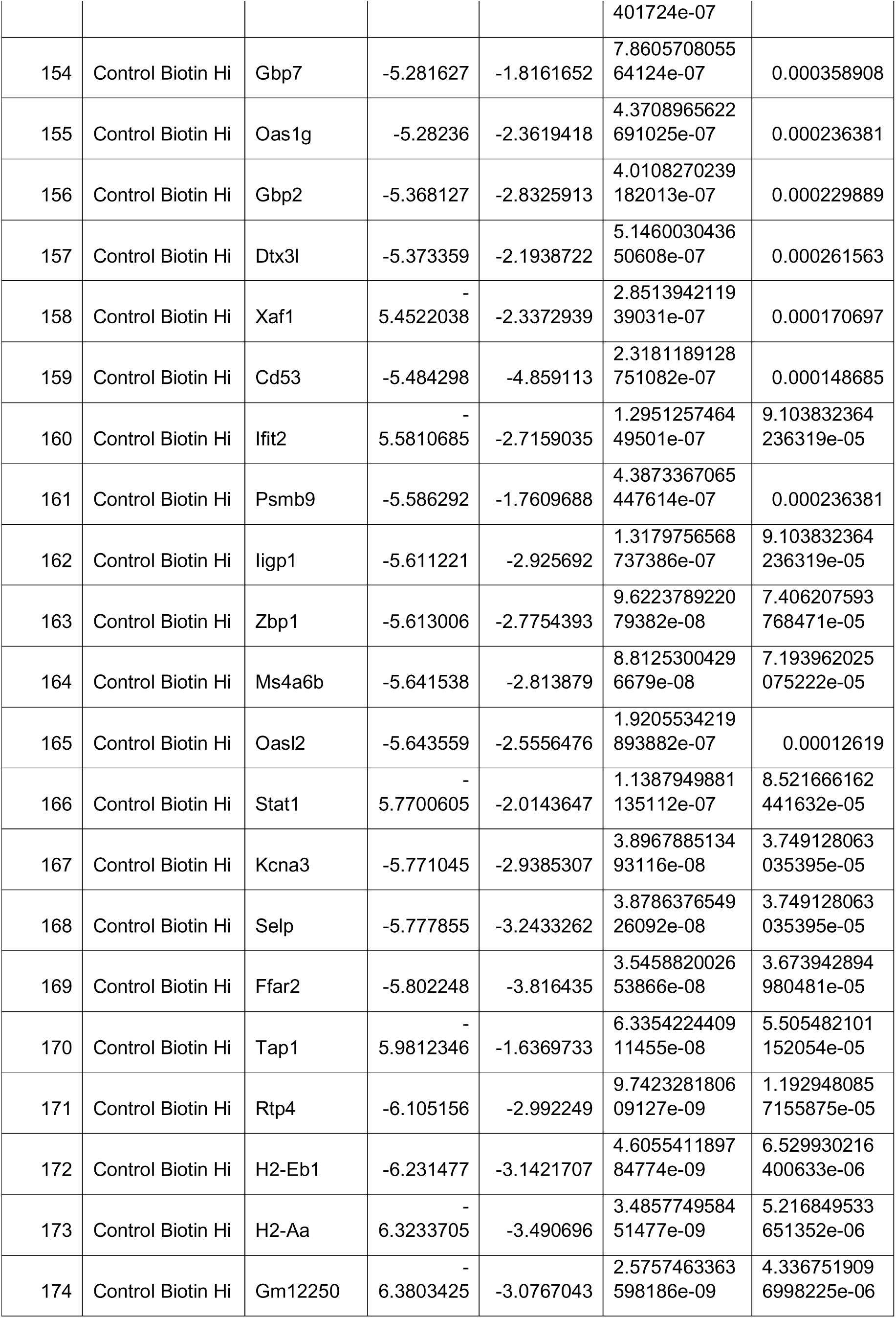
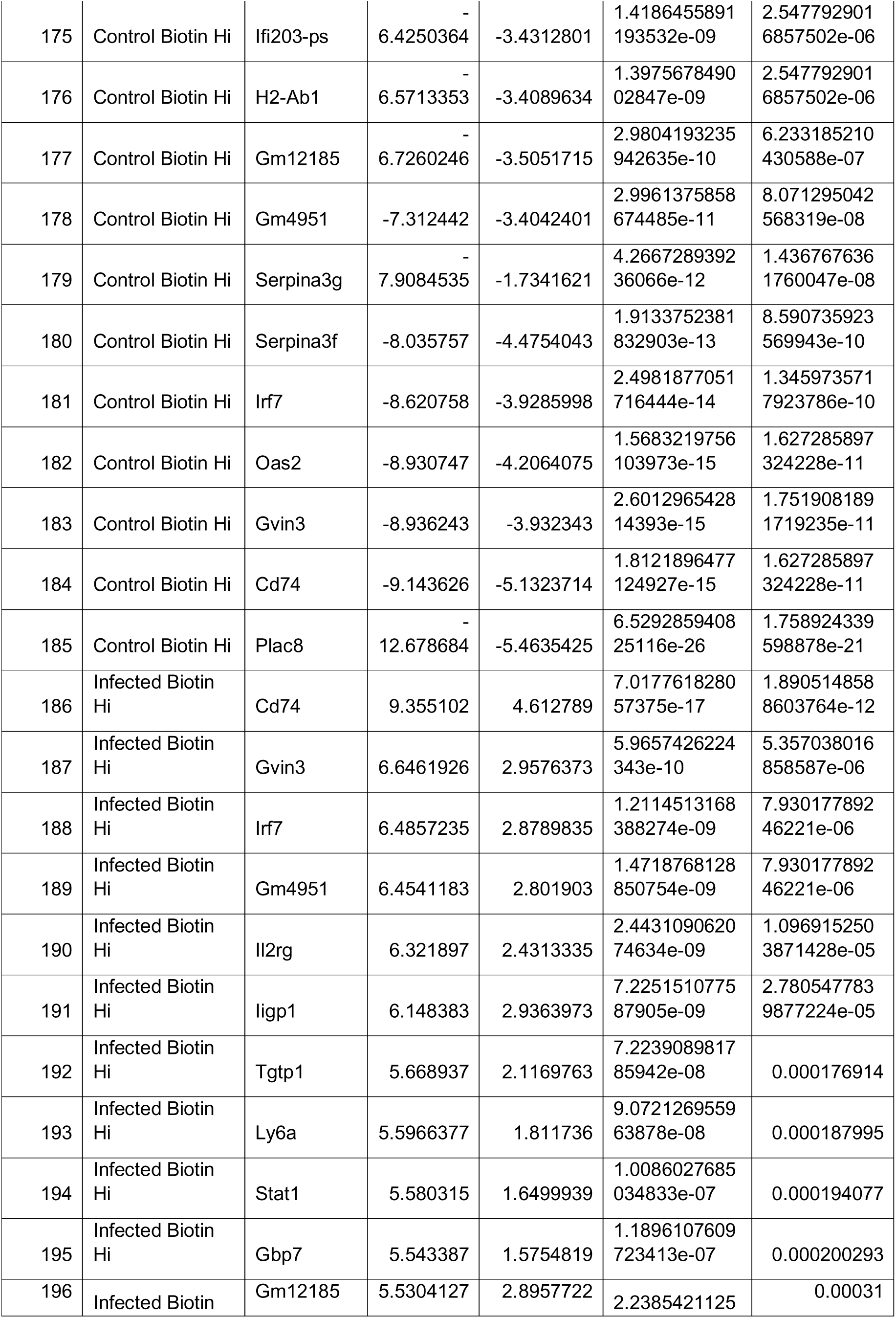
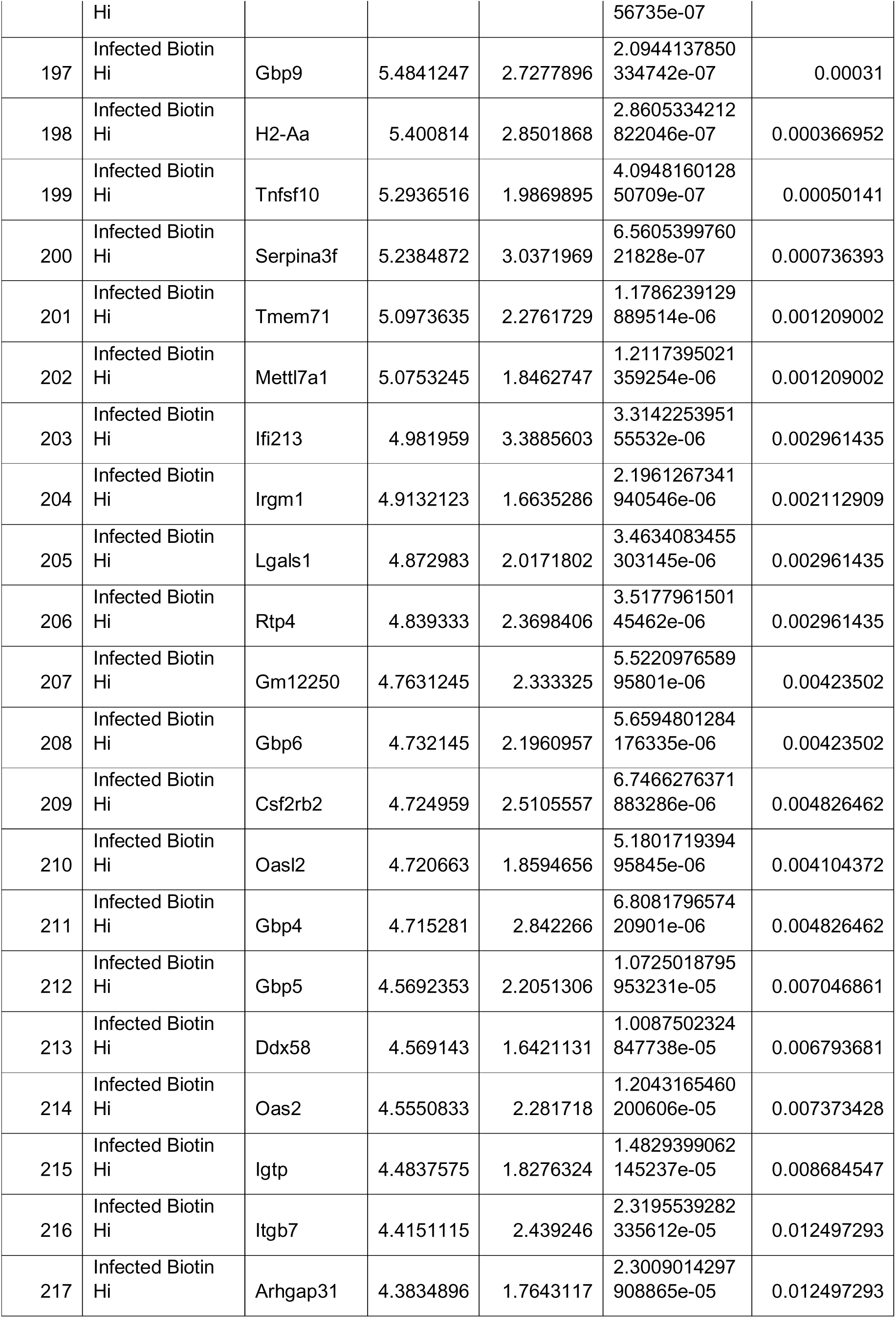
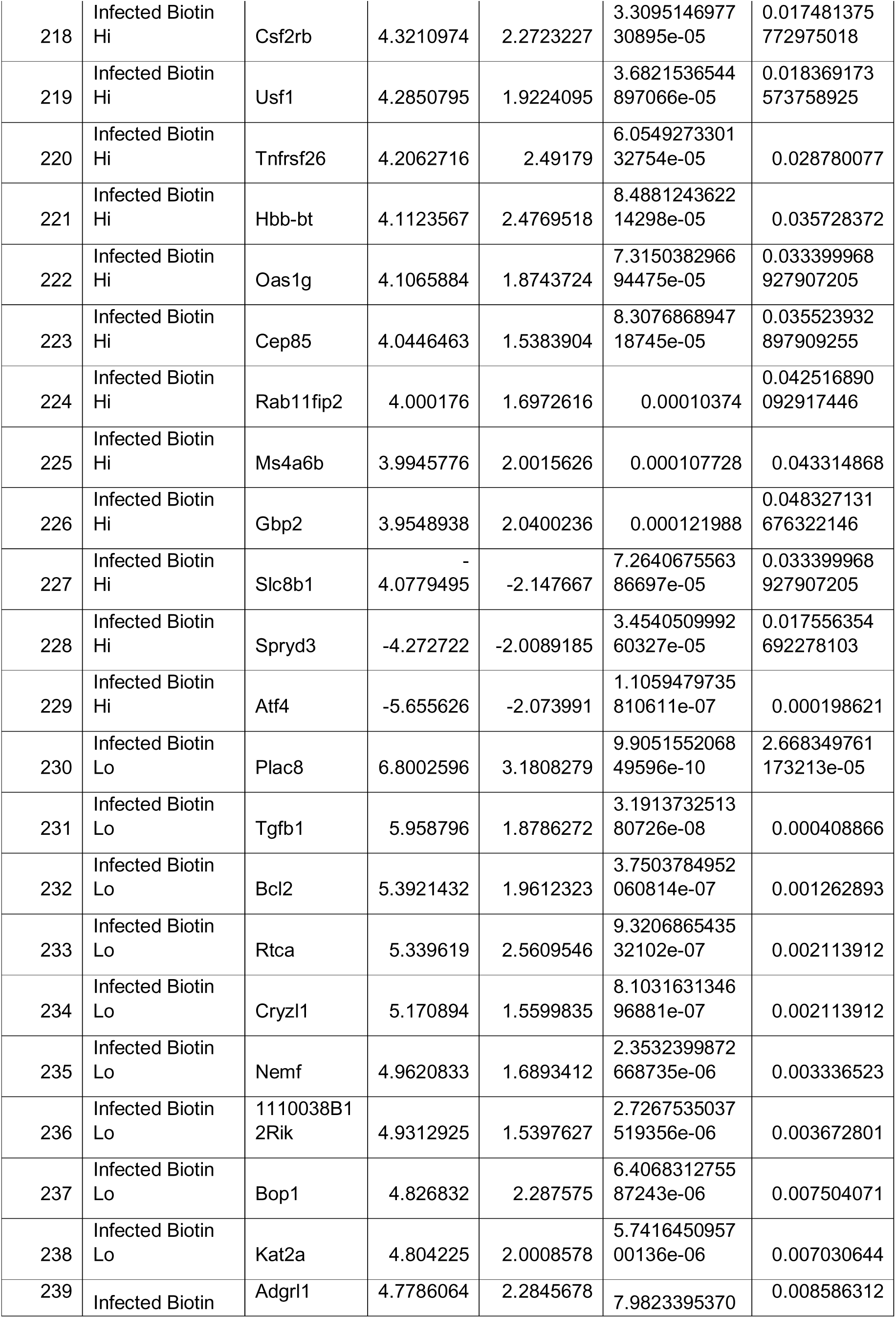
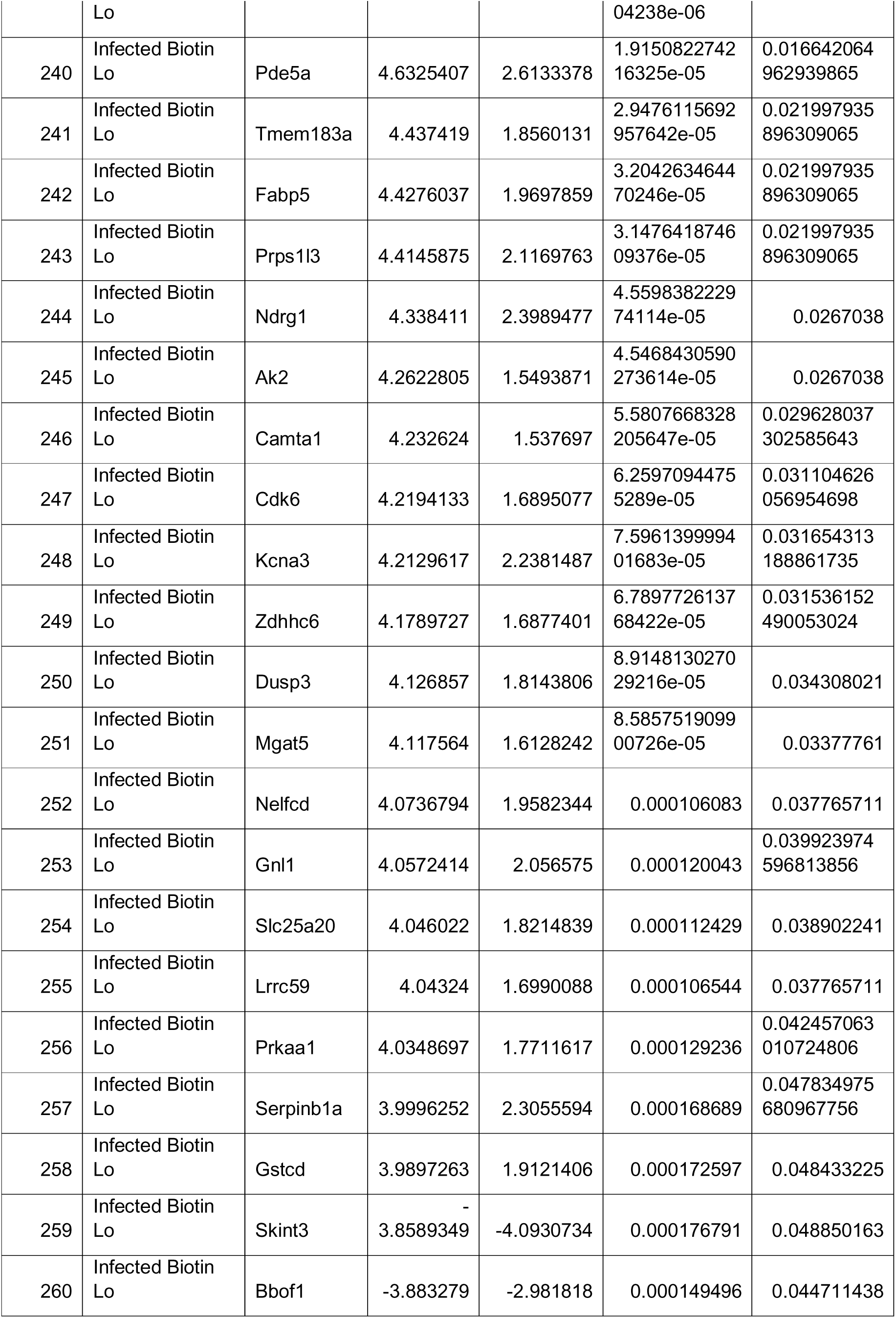
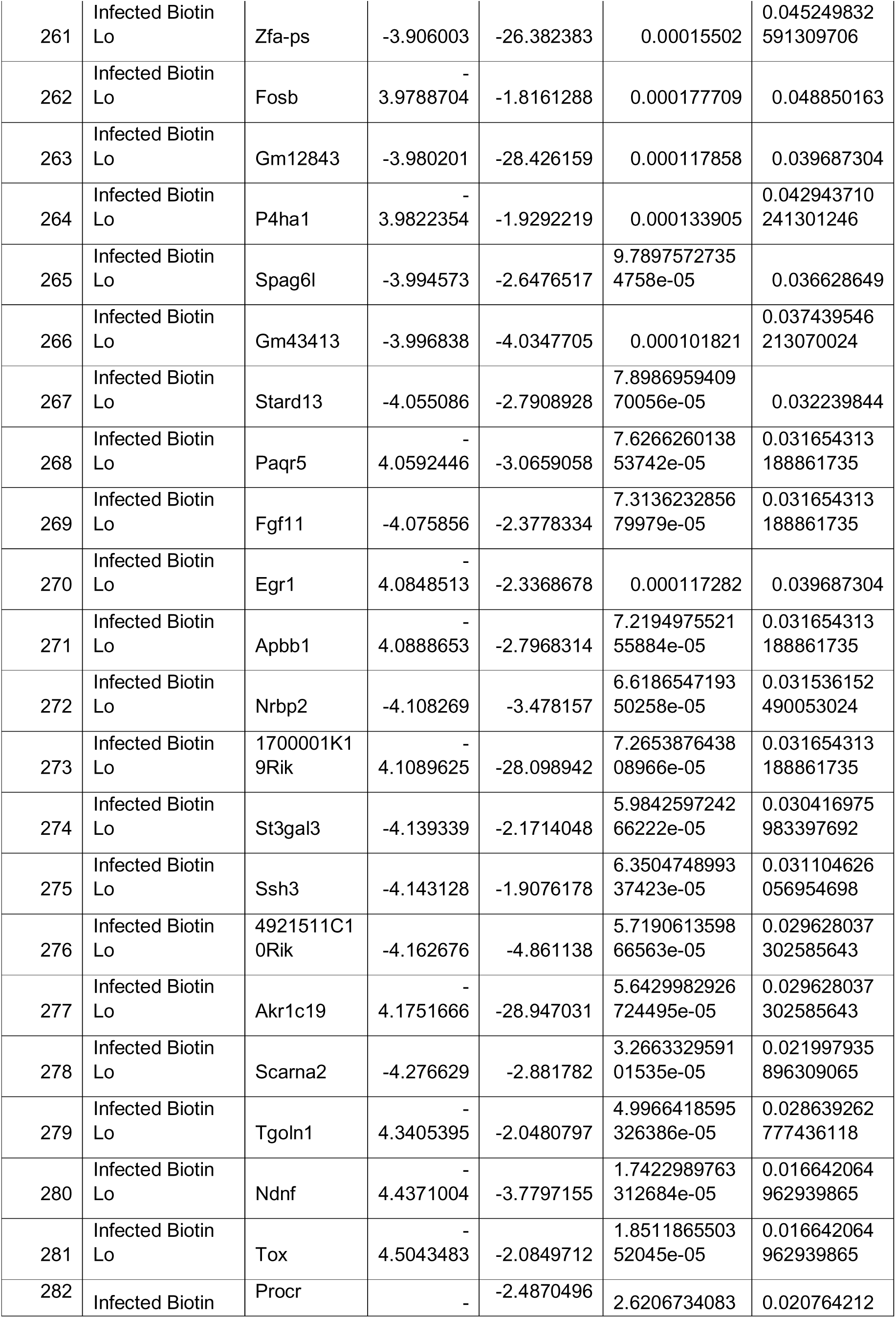
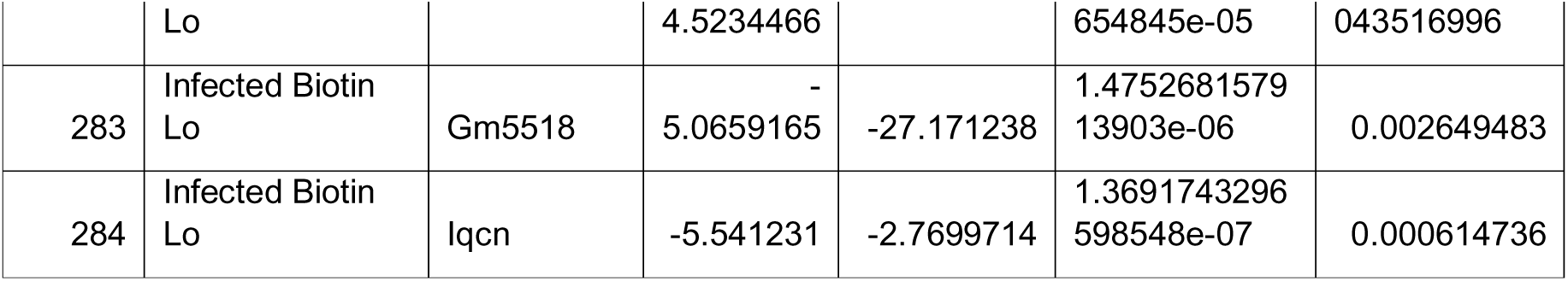
DEGs unique to each cell group, identified through single cell transcriptomics analysis.

**Supplementary Table 2.**
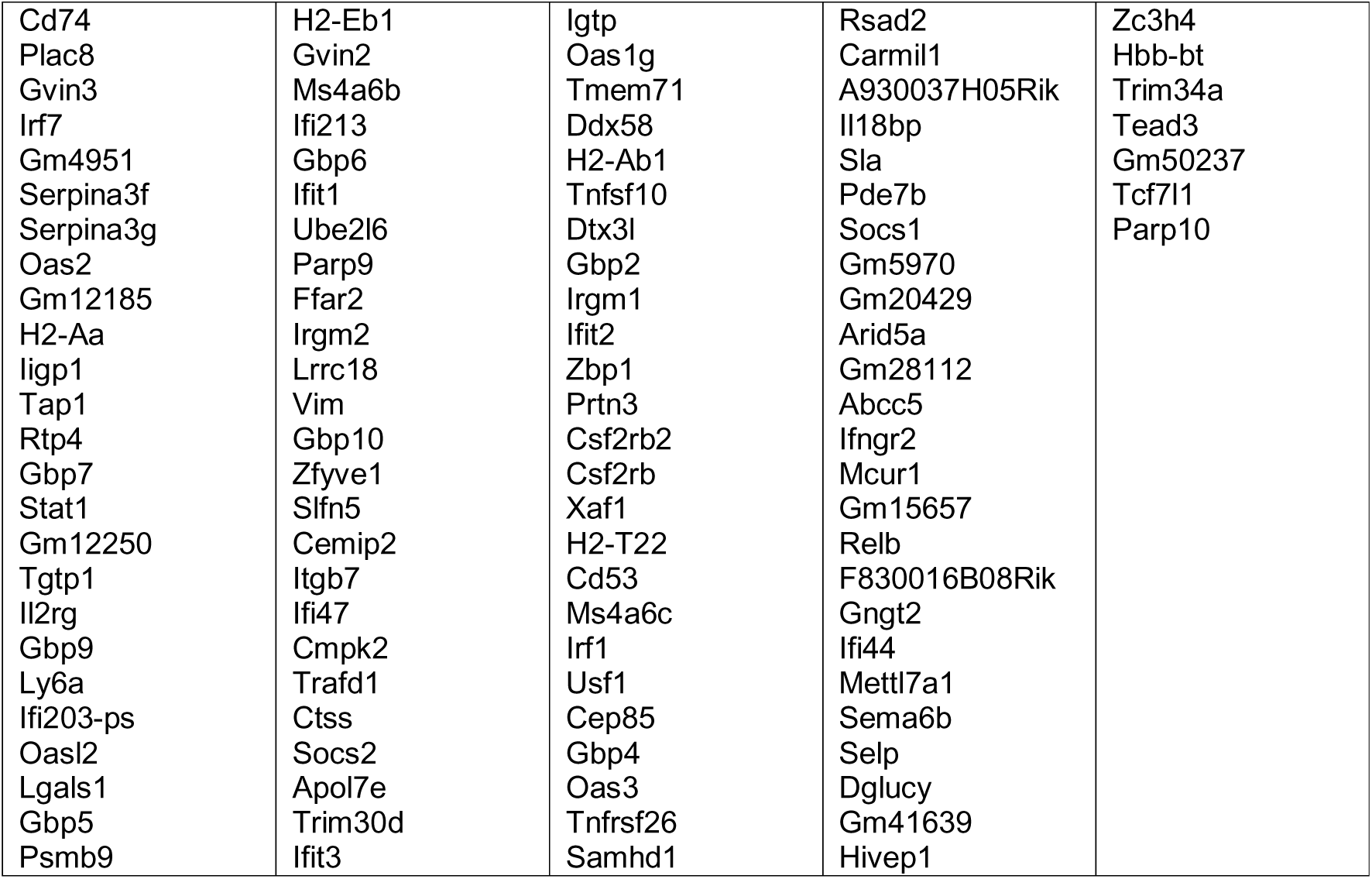
DEGs with significant adjusted p values between Biotin^Hi^ control and Biotin^Hi^ infected HSCs.

**Supplementary Table 3.** Antibodies, reagents, equipment and software used in this study.

| Primary Transplant/Index Sort experiments/FACS Experiments | Catalogue Number | Company | Clone |
| --- | --- | --- | --- |
| FITC anti-mouse/human CD45R/B220 Antibody | 103206 | Biolegend | RA3-6B2 |
| FITC anti-mouse CD4 Antibody | 100510 | Biolegend | RM4-5 |
| FITC anti-mouse CD5 Antibody | 100606 | Biolegend | 53.7-3 |
| FITC anti-mouse CD8a Antibody | 100706 | Biolegend | 53-6.7 |
| FITC anti-mouse Ly-6G/Ly-6C (Gr-1) Antibody | 108406 | Biolegend | RB6-8C5 |
| FITC anti-mouse TER-119/Erythroid Cells Antibody | 116206 | Biolegend | TER-119 |
| Brilliant Violet 421™ Streptavidin | 405226 | Biolegend |  |
| PE/Cyanine7 anti-mouse CD48 [Clone: HM48-1] 100 µg | 103424 | Biolegend | HM48-1 |
| Brilliant Violet 650™ anti-mouse CD150 (SLAM) Antibody | 115931 | Biolegend | TC15-12F12.2 |
| Sca-1PerCP Cy5.5 anti-mouse Ly6A/E (Sca1) | 108124 | Biolegend | D7 |
| Anti-Mouse EPCR Antibody, Clone RMEPCR1560 (1560) | 60038PE | Stem Cell Technologies | RMEPCR1560 (1560) |
| APC/Cyanine7 anti-mouse CD117 (c-kit) [Clone: 2B8] 100 microg | 105826 | Biolegend | 2B8 |
| Brilliant Violet 785™ anti-mouse CD45 Antibody | 103149 | Biolegend | 30-F11 |
| CD45 BUV395 30-F11 50ug | 564279 | BD | 30-F11 |
| BUV395 anti-mouse CD4 Antibody | 740208 | BD | RM4-5 |
| BUV395anti-mouse/human CD45R/B220 Antibody | 563793 | BD | RA3-6B2 |
| BUV395 anti-mouse TER-119/Erythroid Cells Antibody | 563827 | BD | TER-119 |
| BUV395 anti-mouse CD5 Antibody | 740206 | BD | 53.7-3 |
| BUV395 anti-mouse CD8a Antibody | 563786 | BD | 53-6.7 |
| BUV395 anti-mouse Ly-6G/Ly-6C (Gr-1) Antibody | 563849 | BD | RB6-8C5 |
| Brilliant Violet 711™ anti-mouse CD45 [Clone: 30-F11] 50 microg | 103147 | Biolegend | 30-F11 |
| Fixable Viability stain 510 | 564406 | BD Biosciences |  |
| APC anti-mouse CD135 Antibody | 135310 | Biolegend | A2F10 |
| Brilliant Violet 421™ Streptavidin | 405225 | Biolegend |  |
| Fixable Viability Stain 780 | 565388 | BD Horizon |  |
| Brilliant Violet 510™ anti-mouse/human CD45R/B220 Antibody | 103247 | Biolegend | RA3-6B2 |
| Brilliant Violet 510™ anti-mouse TER-119/Erythroid Cells Antibody | 116237 | Biolegend | 116237 |
| Brilliant Violet 510™ anti-mouse Ly-6G/Ly-6C (Gr-1) Antibody | 108437 | Biolegend | 108437 |
| Brilliant Violet 510™ anti-mouse CD4 Antibody | 100553 | Biolegend | 100553 |
| Brilliant Violet 510™ anti-mouse CD5 Antibody | 100627 | Biolegend | 100627 |
| Brilliant Violet 510™ anti-mouse CD8a Antibody | 100751 | Biolegend | 53-6.7 |
| CD201 (EPCR) Monoclonal Antibody (eBio1560 (1560)), APC, eBioscience | 17-20120-82 | eBioscience | 1560 |
| CD201 (EPCR) Monoclonal Antibody (eBio1560 (1560)), eFluor 710 eBioscience | 46-2012-80 | eBioscience | 1560 |
| CD144 (VE-cadherin) Monoclonal Antibody (eBioBV13 (BV13)), eFluor 660, eBioscience™ | 50-1441-82 | ThermoFisher | 50-1441-82 |
| CD117 Microbeads, mouse | 130-091-224 | Miltenyi |  |
| LS Columns | 130-042-401 | Miltenyi |  |
| OneComp eBeads Compensation Beads | 01-1111-42 | Biolegend |  |
| Brilliant stain Buffer | 563794 | BD Biosciences |  |
| Ki67 | 564071 | BD Biosciences |  |
| SYTOX Green | M36008 | Thermofisher |  |
| Multilineage Reconstitution |  |  |  |
| PE/Cyanine7 anti-mouse CD45.1 Antibody | 110730 | Biolegend | A20 |
| FITC anti-mouse CD45.2 Antibody | 109806 | Biolegend | 104 |
| APC anti-mouse CD3ε Antibody | 100312 | Biolegend | 100312 |
| Pacific Blue™ anti-mouse CD19 Antibody | 115523 | Biolegend | 6D5 |
| PE anti-mouse/human CD11b Antibody | 101208 | Biolegend | M1/70 |
| Viability 780 | 565388 | BD Horizon |  |
| BM Stroma FACS |  |  |  |
| DNAseI | 260913-25MU | Calbiochem |  |
| anti-CD45 microbeads, mouse 2ml | 130-052-301 | Miltenyi |  |
| Collagenase Type I - 5 gm | LS004197 | Worthington |  |
| Collagenase Type IV 1-gm | LS004188 | Worthington |  |
| PerCP/Cy5.5 anti-mouse Ly-6A/E (Sca-1) Antibody | 108124 | Biolegend | D7 |
| PE/Cy7 anti-mouse TER-119/Erythroid Cells Antibody | 116221 | Biolegend | TER-119 |
| PE/Cy7 anti-mouse CD31 Antibody | 102524 | Biolegend | MEC13.3 |
| Alexa Fluor® 700 anti-mouse CD45.2 Antibody | 109821 | Biolegend | 104 |
| CD140a (PDGFRA) Monoclonal Antibody (APA5) | 17-1401-81 | Thermofisher | APA5 |
| Mouse Leptin R Biotinylated Antibody | BAF497 | R&D Systems |  |
| BUV395 Rat Anti-Mouse CD51 | 747834 | BD | RMV-7 |
| Mouse ALCAM/CD166 PE-conjugated Antibody | FAB1172P | R&D Systems |  |
| BV421 Streptavidin | 405226 | Biolegend |  |
| single cell and minibulk RNA sequencing |  |  |  |
| SUPERase-In RNase Inhibitor | AM2694 | Ambion |  |
| 10% Triton X-100 | 93443 | Sigma |  |
| DEPC-treated or Rnase-free H2O | 129112 | Qiagen |  |
| Commercial Assays |  |  |  |
| CellROX™ Deep Red Reagent, for oxidative stress detection | C10422 | Thermofisher |  |
| CellROX® Green Reagent | C10444 | Thermofisher |  |
| Click-iT™ EdU Alexa Fluor™ 647 Flow Cytometry Assay Kit | C10424 | Thermofisher |  |
| mitoSOX Red | M36008 | Invitrogen |  |
| MitoTracker Green | M7514 | ThermoFisher |  |
| TMRM Assay Kit for Flow Cytometry | M20036 | Thermofisher |  |
| Chemicals |  |  |  |
| EZ-Link™ Sulfo-NHS-LC-LC-Biotin Thermo Scientific PI21338; 50mg | 21338 | Thermofisher |  |
| Doxycycline | D9891-5G | Sigma |  |
| N-Acetyl-L-cysteine | A7250-5G | Sigma |  |
| EdU (5-ethynyl-2'-deoxyuridine) | A10044 | Thermofisher |  |
| GIEMSA | 48900-500ML-F | Sigma |  |
| Solutions |  |  |  |
| EDTA 0.5M pH8.0 +/-1 (100mL) | 46-034-CI | Corning | 0.5 M stock |
| Phosphate BDulbecco's Phosphate Buffered Saline,Modified, without calcium chloride and magnesium chloride | D8537 | Sigma Aldrich |  |
| Alsever's Solution | A3551 | Sigma Aldrich |  |
| BD Cytofix Cytoperm | 561651 | BD Bioscience |  |
| DMEM | Gibco | 11960-044 |  |
| Drugs |  |  |  |
| Parathyroid Hormone | 4012212 | Bachem | rat PTH 1-34 |
| Phenylhydrazine | P26252 | Sigma |  |
| Atovaquoe | A7986-50MG | Sigma |  |
| Cryosectioning |  |  |  |
| CFSA Cryojane adhesive slides | 39475208 | Leica Biosystems |  |
| Cryojane Adhesive Tape Windows | 39475214 | Leica Biosystems |  |
| Immunostaining |  |  |  |
| Mouse Leptin R Biotinylated Antibody | BAF497 | R&D Systems |  |
| Purified anti-mouse CD117 | 105802 | Biolegend | 2B8 |
| Streptavidin, Alexa Fluor™ 647 conjugate | S-32357 | Thermofisher |  |
| Normal donkey serum | ab7475 | Abcam |  |
| horse serum | 31874 | Life Technologies |  |
| DAPI | D9542 | Thermofisher |  |
| Prolong Antifade Diamond Mountant | P36961 | Thermofisher |  |
| Analysis Softwares used |  |  |  |
| GraphPad Prism |  |  |  |
| Imaris 10.0 |  |  |  |
| Fiji |  |  |  |
| FlowJo |  |  |  |
| Python or R |  |  |  |
| Zen Blue |  |  |  |
| Equipmnet |  |  |  |
| BD Fortessa |  |  |  |
| BS Symphony A5 |  |  |  |
| BD FACS Aria sorter |  |  |  |
| LSN ZEIS 960 confocal microscope |  |  |  |
| Leica Cryostat CM3050S |  |  |  |
| Zeis Bright field microscope |  |  |  |
| Illumina Hiseq4000 |  |  |  |

## Notes

### Competing Interest Statement

The authors have declared no competing interest.

## References

1. Akinduro, O., Weber, T. S., Ang, H., Haltalli, M. L. R., Ruivo, N., Duarte, D., Rashidi, N. M., Hawkins, E. D., Duffy, K. R., & Lo Celso, C. (2018). Proliferation dynamics of acute myeloid leukaemia and haematopoietic progenitors competing for bone marrow space. Nature Communications, 9(1). 10.1038/s41467-017-02376-5

2. Alghamdi, N., Chang, W., Dang, P., Lu, X., Wan, C., Gampala, S., Huang, Z., Wang, J., Ma, Q., Zang, Y., Fishel, M., Cao, S., & Zhang, C. (2021). A graph neural network model to estimate cell-wise metabolic flux using single-cell RNA-seq data. Genome Research, 31(10), 1867–1884. 10.1101/gr.271205.120

3. Ansó, E., Weinberg, S. E., Diebold, L. P., Thompson, B. J., Malinge, S., Schumacker, P. T., Liu, X., Zhang, Y., Shao, Z., Steadman, M., Marsh, K. M., Xu, J., Crispino, J. D., & Chandel, N. S. (2017). The mitochondrial respiratory chain is essential for haematopoietic stem cell function. Nature Cell Biology, 19(6), 614–625. 10.1038/ncb3529

4. Arthur Flohr Svendsen, 1 Daozheng Yang,1 KyungMok Kim,2 Seka Lazare,1 Natalia Skinder,1 Erik Zwart,1 Anna Mura-Meszaros,2 Albertina Ausema,1 Bj€orn von Eyss,2 Gerald de Haan,1,* and Leonid Bystrykh1,*. (2021). A comprehensive transcriptome signature of murine hematopoietic stem cell aging. Blood, 136(19), 2125–2132. 10.1182/BLOOD.2019000962

5. Ashley, E. A., Pyae Phyo, A., & Woodrow, C. J. (2018). Malaria. The Lancet, 391(10130), 1608–1621. 10.1016/S0140-6736(18)30324-6

6. Baldridge, M. T., King, K. Y., Boles, N. C., Weksberg, D. C., & Goodell, M. A. (2010). Quiescent haematopoietic stem cells are activated by IFN-γ in response to chronic infection. Nature, 465(7299), 793–797. 10.1038/nature09135

7. Belyaev, N. N., Biró, J., Langhorne, J., & Potocnik, A. J. (2013). Extramedullary Myelopoiesis in Malaria Depends on Mobilization of Myeloid-Restricted Progenitors by IFN-γ Induced Chemokines. PLoS Pathogens, 9(6). 10.1371/journal.ppat.1003406

8. Bogeska, R., Mikecin, A. M., Kaschutnig, P., Fawaz, M., Büchler-Schäff, M., Le, D., Ganuza, M., Vollmer, A., Paffenholz, S. V., Asada, N., Rodriguez-Correa, E., Frauhammer, F., Buettner, F., Ball, M., Knoch, J., Stäble, S., Walter, D., Petri, A., Carreño-Gonzalez, M. J., … Milsom, M. D. (2022a). Inflammatory exposure drives long-lived impairment of hematopoietic stem cell self-renewal activity and accelerated aging. Cell Stem Cell, 29(8), 1273–1284.e8. 10.1016/j.stem.2022.06.012

9. Bowling, S., Sritharan, D., Osorio, F. G., Nguyen, M., Cheung, P., Rodriguez-Fraticelli, A., Patel, S., Yuan, W. C., Fujiwara, Y., Li, B. E., Orkin, S. H., Hormoz, S., & Camargo, F. D. (2020). An Engineered CRISPR-Cas9 Mouse Line for Simultaneous Readout of Lineage Histories and Gene Expression Profiles in Single Cells. Cell, 181(6), 1410–1422.e27. 10.1016/j.cell.2020.04.048

10. Bujanover, N., Goldstein, O., Greenshpan, Y., Turgeman, H., & Klainberger, A. (2020). Identi fi cation of immune-activated hematopoietic stem cells. 133(mC), 2016–2020.

11. Cabezas-Wallscheid, N., Buettner, F., Sommerkamp, P., Klimmeck, D., Ladel, L., Thalheimer, F. B., Pastor-Flores, D., Roma, L. P., Renders, S., Zeisberger, P., Przybylla, A., Schönberger, K., Scognamiglio, R., Altamura, S., Florian, C. M., Fawaz, M., Vonficht, D., Tesio, M., Collier, P., … Trumpp, A. (2017). Vitamin A-Retinoic Acid Signaling Regulates Hematopoietic Stem Cell Dormancy. Cell, 169(5), 807–823.e19. 10.1016/j.cell.2017.04.018

12. Cai, Z., Kotzin, J. J., Ramdas, B., Chen, S., Nelanuthala, S., Palam, L. R., Pandey, R., Mali, R. S., Liu, Y., Kelley, M. R., Sandusky, G., Mohseni, M., Williams, A., Henao-Mejia, J., & Kapur, R. (2018). Inhibition of Inflammatory Signaling in Tet2 Mutant Preleukemic Cells Mitigates Stress-Induced Abnormalities and Clonal Hematopoiesis. Cell Stem Cell, 23(6), 833–849.e5. 10.1016/j.stem.2018.10.013

13. Carrelha, J., Meng, Y., Kettyle, L. M., Luis, T. C., Norfo, R., Alcolea, V., Boukarabila, H., Grasso, F., Gambardella, A., Grover, A., Högstrand, K., Lord, A. M., Sanjuan-Pla, A., Woll, P. S., Nerlov, C., & Jacobsen, S. E. W. (2018). Hierarchically related lineage-restricted fates of multipotent haematopoietic stem cells. Nature, 554(7690), 106–111. 10.1038/nature25455

14. Che, J. L. C., Bode, D., Kucinski, I., Cull, A. H., Bain, F., Becker, H. J., Jassinskaja, M., Barile, M., Boyd, G., Belmonte, M., Zeng, A. G. X., Igarashi, K. J., Rubio-Lara, J., Shepherd, M. S., Clay, A., Dick, J. E., Wilkinson, A. C., Nakauchi, H., Yamazaki, S., … Kent, D. G. (2022a). Identification and characterization of in vitro expanded hematopoietic stem cells. EMBO Reports, 23(10), 1–19. 10.15252/embr.202255502

15. Coban, C., Lee, M. S. J., & Ishii, K. J. (2018). Tissue-specific immunopathology during malaria infection. Nature Reviews Immunology, 18(4), 266–278. 10.1038/nri.2017.138

16. de Almeida, M. J., Luchsinger, L. L., Corrigan, D. J., Williams, L. J., & Snoeck, H. W. (2017). Dye-Independent Methods Reveal Elevated Mitochondrial Mass in Hematopoietic Stem Cells. Cell Stem Cell, 21(6), 725–729.e4. 10.1016/j.stem.2017.11.002

17. Donato, E., Correia, N., Andresen, C., Karpova, D., Würth, R., Klein, C., Sohn, M., Przybylla, A., Zeisberger, P., Rothfelder, K., Salih, H., Bonig, H., Stasik, S., Röllig, C., Dolnik, A., Bullinger, L., Buchholz, F., Thiede, C., Hübschmann, D., & Trumpp, A. (2023). Retained functional normal and preleukemic HSCs at diagnosis are associated with good prognosis in DNMT3AmutNPM1mut AMLs. Blood Advances, 7(6), 1011–1018. 10.1182/bloodadvances.2022008497

18. Dykstra, B., Kent, D., Bowie, M., McCaffrey, L., Hamilton, M., Lyons, K., Lee, S. J., Brinkman, R., & Eaves, C. (2007). Long-Term Propagation of Distinct Hematopoietic Differentiation Programs In Vivo. Cell Stem Cell, 1(2), 218–229. 10.1016/j.stem.2007.05.015

19. Essers, M. A. G., Offner, S., Blanco-Bose, W. E., Waibler, Z., Kalinke, U., Duchosal, M. A., & Trumpp, A. (2009). IFNα activates dormant haematopoietic stem cells in vivo. Nature, 458(7240), 904– 908. 10.1038/nature07815

20. Fernández-Vizarra, E., López-Calcerrada, S., Sierra-Magro, A., Pérez-Pérez, R., Formosa, L. E., Hock, D. H., Illescas, M., Peñas, A., Brischigliaro, M., Ding, S., Fearnley, I. M., Tzoulis, C., Pitceathly, R. D. S., Arenas, J., Martín, M. A., Stroud, D. A., Zeviani, M., Ryan, M. T., & Ugalde, C. (2022). Two independent respiratory chains adapt OXPHOS performance to glycolytic switch. Cell Metabolism, 34(11), 1792–1808.e6. 10.1016/j.cmet.2022.09.005

21. Filippi, M. D., & Ghaffari, S. (2019). Mitochondria in the maintenance of hematopoietic stem cells: New perspectives and opportunities. Blood, 133(18), 1943–1952. 10.1182/blood-2018-10-808873

22. Haltalli, M. L. R., Watcham, S., Wilson, N. K., Eilers, K., Ang, H., Birch, F., Anton, S. G., Pirillo, C., Ruivo, N., Vainieri, M. L., Pospori, C., Sinden, R. E., Luis, T. C., Duffy, K. R., Blagborough, A. M., & Lo, C. (2020). Manipulating niche composition limits damage to haematopoietic stem cells during Plasmodium infection.22. 1–47.

23. Hamey, F. K., & Göttgens, B. (2019). Machine learning predicts putative hematopoietic stem cells within large single-cell transcriptomics data sets. Experimental Hematology, 78, 11–20. 10.1016/j.exphem.2019.08.009

24. Hernández-Malmierca, P., Vonficht, D., Schnell, A., Uckelmann, H. J., Bollhagen, A., Mahmoud, M. A. A., Landua, S. L., van der Salm, E., Trautmann, C. L., Raffel, S., Grünschläger, F., Lutz, R., Ghosh, M., Renders, S., Correia, N., Donato, E., Dixon, K. O., Hirche, C., Andresen, C., … Haas, S. (2022). Antigen presentation safeguards the integrity of the hematopoietic stem cell pool. Cell Stem Cell, 29(5), 760–775.e10. 10.1016/j.stem.2022.04.007

25. Hinge, A., He, J., Bartram, J., Javier, J., Xu, J., Fjellman, E., Sesaki, H., Li, T., Yu, J., Wunderlich, M., Mulloy, J., Kofron, M., Salomonis, N., Grimes, H. L., & Filippi, M. D. (2020). Asymmetrically Segregated Mitochondria Provide Cellular Memory of Hematopoietic Stem Cell Replicative History and Drive HSC Attrition. Cell Stem Cell, 26(3), 420–430.e6. 10.1016/j.stem.2020.01.016

26. Hirche, C., Frenz, T., Haas, S. F., Döring, M., Borst, K., Tegtmeyer, P. K., Brizic, I., Jordan, S., Keyser, K., Chhatbar, C., Pronk, E., Lin, S., Messerle, M., Jonjic, S., Falk, C. S., Trumpp, A., Essers, M. A. G., & Kalinke, U. (2017). Systemic Virus Infections Differentially Modulate Cell Cycle State and Functionality of Long-Term Hematopoietic Stem Cells In Vivo. Cell Reports, 19(11), 2345–2356. 10.1016/j.celrep.2017.05.063

27. Hormaechea-Agulla, D., Matatall, K. A., Le, D. T., Kain, B., Long, X., Kus, P., Jaksik, R., Challen, G. A., Kimmel, M., & King, K. Y. (2021). Chronic infection drives Dnmt3a-loss-of-function clonal hematopoiesis via IFNγ signaling. Cell Stem Cell, 28(8), 1428–1442.e6. 10.1016/j.stem.2021.03.002

28. Isobe, T., Kucinski, I., Barile, M., Wang, X., Hannah, R., Bastos, H. P., Chabra, S., Vijayabaskar, M. S., Sturgess, K. H. M., Williams, M. J., Giotopoulos, G., Marando, L., Li, J., Rak, J., Gozdecka, M., Prins, D., Shepherd, M. S., Watcham, S., Green, A. R., … Göttgens, B. (2023). Preleukemic single-cell landscapes reveal mutation-specific mechanisms and gene programs predictive of AML patient outcomes. Cell Genomics, 3(12). 10.1016/j.xgen.2023.100426

29. Karpova, D., Huerga Encabo, H., Donato, E., Kotova, I., Calderazzo, S., Leppä, AM., Panten, J., Przbylla, A., Seifried, E., Kopp-Schneider, A., Wong, TN., Bonnet, D., Bonig, H., & Trumpp, A. (2025). Clonal Hematopoiesis Landscape in Frequent Blood Donors. 10.1101/2022.07.24.22277825

30. Kaufmann, K. B., Zeng, A. G. X., Coyaud, E., Garcia-Prat, L., Papalexi, E., Murison, A., Laurent, E. M. N., Chan-Seng-Yue, M., Gan, O. I., Pan, K., McLeod, J., Boutzen, H., Zandi, S., Takayanagi, S. ichiro, Satija, R., Raught, B., Xie, S. Z., & Dick, J. E. (2021). A latent subset of human hematopoietic stem cells resists regenerative stress to preserve stemness. Nature Immunology, 22(6), 723–734. 10.1038/s41590-021-00925-1

31. Kent, D. G., Copley, M. R., Benz, C., Wöhrer, S., Dykstra, B. J., Ma, E., Cheyne, J., Zhao, Y., Bowie, M. B., Zhao, Y., Gasparetto, M., Delaney, A., Smith, C., Marra, M., & Eaves, C. J. (2009). Prospective isolation and molecular characterization of hematopoietic stem cells with durable self-renewal potential. Blood, 113(25), 6342–6350. 10.1182/blood-2008-12-192054

32. King, K. Y., Baldridge, M. T., Weksberg, D. C., Chambers, S. M., Lukov, G. L., Wu, S., Boles, N. C., Jung, S. Y., Qin, J., Liu, D., Songyang, Z., Eissa, N. T., Taylor, G. A., & Goodell, M. A. (2011). Irgm1 protects hematopoietic stem cells by negative regulation of IFN signaling. Blood, 118(6), 1525–1533. 10.1182/blood-2011-01-328682

33. Kowalczyk, M. S., Tirosh, I., Heckl, D., Rao, T. N., Dixit, A., Haas, B. J., Schneider, R. K., Wagers, A. J., Ebert, B. L., & Regev, A. (2015a). Single-cell RNA-seq reveals changes in cell cycle and differentiation programs upon aging of hematopoietic stem cells. Genome Research, 25(12), 1860–1872. 10.1101/gr.192237.115

34. Kucinski, I., Campos, J., Barile, M., Severi, F., Bohin, N., Moreira, P. N., Allen, L., Lawson, H., Haltalli, M. L. R., Kinston, S. J., O’Carroll, D., Kranc, K. R., & Göttgens, B. (2024). A time– and single-cell-resolved model of murine bone marrow hematopoiesis. Cell Stem Cell, 31(2), 244–259.e10. 10.1016/j.stem.2023.12.001

35. Kwak, M., & Ghazizadeh, S. (2015). Analysis of histone H2BGFP retention in mouse submandibular gland reveals actively dividing stem cell populations. Stem Cells and Development, 24(5), 565–574. 10.1089/scd.2014.0355

36. Laurenti, E., Frelin, C., Xie, S., Ferrari, R., Dunant, C. F., Zandi, S., Neumann, A., Plumb, I., Doulatov, S., Chen, J., April, C., Fan, J. B., Iscove, N., & Dick, J. E. (2015). CDK6 levels regulate quiescence exit in human hematopoietic stem cells. Cell Stem Cell, 16(3), 302–313. 10.1016/j.stem.2015.01.017

37. Li, J., Williams, M. J., Park, H. J., Bastos, H. P., Wang, X., Prins, D., Wilson, N. K., Johnson, C., Sham, K., Wantoch, M., Watcham, S., Kinston, S. J., Pask, D. C., Hamilton, T. L., Sneade, R., Waller, A. K., Ghevaert, C., Vassiliou, G. S., Laurenti, E., … Green, A. R. (2022). STAT1 is essential for HSC function and maintains MHCIIhi stem cells that resist myeloablation and neoplastic expansion. Blood, 140(14), 1592–1606. 10.1182/blood.2021014009

38. Liang, R., Arif, T., Kalmykova, S., Kasianov, A., Lin, M., Menon, V., Qiu, J., Bernitz, J. M., Moore, K., Lin, F., Benson, D. L., Tzavaras, N., Mahajan, M., Papatsenko, D., & Ghaffari, S. (2020). Restraining Lysosomal Activity Preserves Hematopoietic Stem Cell Quiescence and Potency. Cell Stem Cell, 26(3), 359–376.e7. 10.1016/j.stem.2020.01.013

39. Luis, T. C., Barkas, N., Carrelha, J., Giustacchini, A., Mazzi, S., Norfo, R., Wu, B., Aliouat, A., Guerrero, J. A., Rodriguez-Meira, A., Bouriez-Jones, T., Macaulay, I. C., Jasztal, M., Zhu, G., Ni, H., Robson, M. J., Blakely, R. D., Mead, A. J., Nerlov, C., … Jacobsen, S. E. W. (2023). Perivascular niche cells sense thrombocytopenia and activate hematopoietic stem cells in an IL-1 dependent manner. Nature Communications, 14(1). 10.1038/s41467-023-41691-y

40. Mansell, E., Sigurdsson, V., Deltcheva, E., Brown, J., James, C., Miharada, K., Soneji, S., Larsson, J., & Enver, T. (2021). Mitochondrial Potentiation Ameliorates Age-Related Heterogeneity in Hematopoietic Stem Cell Function. Cell Stem Cell, 28(2), 241–256.e6. 10.1016/j.stem.2020.09.018

41. Matatall, K. A., Jeong, M., Chen, S., Sun, D., Chen, F., Mo, Q., Kimmel, M., & King, K. Y. (2016). Chronic Infection Depletes Hematopoietic Stem Cells through Stress-Induced Terminal Differentiation. Cell Reports, 17(10), 2584–2595. 10.1016/j.celrep.2016.11.031

42. Min, I. M., Pietramaggiori, G., Kim, F. S., Passegué, E., Stevenson, K. E., & Wagers, A. J. (2008). The Transcription Factor EGR1 Controls Both the Proliferation and Localization of Hematopoietic Stem Cells. Cell Stem Cell, 2(4), 380–391. 10.1016/j.stem.2008.01.015

43. Munz, C. M., Dressel, N., Chen, M., Grinenko, T., Roers, A., & Gerbaulet, A. (2023). Regeneration after blood loss and acute inflammation proceeds without contribution of primitive HSCs. Blood, 141(20), 2483–2492. 10.1182/blood.2022018996

44. Muzumdar, M. D., Tasic, B., Miyamichi, K., Li, N., & Luo, L. (2007). A global double-fluorescent cre reporter mouse. Genesis (United States), 45(9), 593–605. 10.1002/dvg.20335

45. Nestorowa, S., Hamey, F. K., Pijuan Sala, B., Diamanti, E., Shepherd, M., Laurenti, E., Wilson, N. K., Kent, D. G., & Göttgens, B. (2016). A single-cell resolution map of mouse hematopoietic stem and progenitor cell differentiation. Blood, 128(8), e20–e31. 10.1182/blood-2016-05-716480

46. Ning, S., Pagano, J. S., & Barber, G. N. (2011). IRF7: Activation, regulation, modification and function. Genes and Immunity, 12(6), 399–414. 10.1038/gene.2011.21

47. Nygren, J. M., & Bryder, D. (2008a). A novel assay to trace proliferation history in vivo reveals that enhanced divisional kinetics accompany loss of hematopoietic stem cell self-renewal. PLoS ONE, 3(11). 10.1371/journal.pone.0003710

48. Papa, L., Djedaini, M., & Hoffman, R. (2019). Mitochondrial role in stemness and differentiation of hematopoietic stem cells. Stem Cells International, 2019. 10.1155/2019/4067162

49. Picelli, S., Björklund, Å. K., Faridani, O. R., Sagasser, S., Winberg, G., & Sandberg, R. (2013). Smart-seq2 for sensitive full-length transcriptome profiling in single cells. Nature Methods, 10(11), 1096–1100. 10.1038/nmeth.2639

50. Pietras, E. M., Mirantes-Barbeito, C., Fong, S., Loeffler, D., Kovtonyuk, L. V., Zhang, S., Lakshminarasimhan, R., Chin, C. P., Techner, J. M., Will, B., Nerlov, C., Steidl, U., Manz, M. G., Schroeder, T., & Passegué, E. (2016). Chronic interleukin-1 exposure drives haematopoietic stem cells towards precocious myeloid differentiation at the expense of self-renewal. Nature Cell Biology, 18(6), 607–618. 10.1038/ncb3346

51. Rodriguez-Fraticelli, A. E., Wolock, S. L., Weinreb, C. S., Panero, R., Patel, S. H., Jankovic, M., Sun, J., Calogero, R. A., Klein, A. M., & Camargo, F. D. (2018). Clonal analysis of lineage fate in native haematopoiesis. Nature, 553(7687), 212–216. 10.1038/nature25168

52. Samarasekera, U. (2023). Climate change and malaria: predictions becoming reality. *Lancet (London*, England), 402(10399), 361–362. 10.1016/S0140-6736(23)01569-6

53. Scheicher, R., Hoelbl-Kovacic, A., Bellutti, F., Tigan, A.-S., Prchal-Murphy, M., Heller, G., Schneckenleithner, C., Salazar-Roa, M., Ochbauer-M, S. Z., Zuber, J., Malumbres, M., Kollmann, K., & Sexl, V. (2015). CDK6 as a key regulator of hematopoietic and leukemic stem cell activation. 10.1182/blood-2014-06

54. Schulte, R., Wilson, N. K., Prick, J. C. M., Cossetti, C., Maj, M. K., Gottgens, B., & Kent, D. G. (2015). Index sorting resolves heterogeneous murine hematopoietic stem cell populations. Experimental Hematology, 43(9), 803–811. 10.1016/j.exphem.2015.05.006

55. Thapa, D., Huang, S. B., Muñoz, A. R., Yang, X., Bedolla, R. G., Hung, C. N., Chen, C. L., Huang, T. H. M., Liss, M. A., Reddick, R. L., Miyamoto, H., Kumar, A. P., & Ghosh, R. (2020). Attenuation of NAD[P]H:quinone oxidoreductase 1 aggravates prostate cancer and tumor cell plasticity through enhanced TGFβ signaling. Communications Biology, 3(1), 1–12. 10.1038/s42003-019-0720-z

56. Vainieri, M. L., Blagborough, A. M., MacLean, A. L., Haltalli, M. L. R., Ruivo, N., Fletcher, H. A., Stumpf, M. P. H., Sinden, R. E., & Celso, C. Lo. (2016a). Systematic tracking of altered haematopoiesis during sporozoitemediated malaria development reveals multiple response points. Open Biology, 6(6). 10.1098/rsob.160038

57. Vannini, N., Girotra, M., Naveiras, O., Nikitin, G., Campos, V., Giger, S., Roch, A., Auwerx, J., & Lutolf, M. P. (2016). Specification of haematopoietic stem cell fate via modulation of mitochondrial activity. Nature Communications, 7, 1–9. 10.1038/ncomms13125

58. Venugopal, K., Hentzschel, F., Valkiūnas, G., & Marti, M. (2020). Plasmodium asexual growth and sexual development in the haematopoietic niche of the host. Nature Reviews Microbiology, 18(3), 177–189. 10.1038/s41579-019-0306-2

59. Wang, M., Brandt, L. T. L., Wang, X., Russell, H., Mitchell, E., Kamimae-Lanning, A. N., Brown, J. M., Dingler, F. A., Garaycoechea, J. I., Isobe, T., Kinston, S. J., Gu, M., Vassiliou, G. S., Wilson, N. K., Göttgens, B., & Patel, K. J. (2023). Genotoxic aldehyde stress prematurely ages hematopoietic stem cells in a p53-driven manner. Molecular Cell, 83(14), 2417–2433.e7. 10.1016/j.molcel.2023.05.035

60. Wilson, N. K., Kent, D. G., Buettner, F., Shehata, M., Macaulay, I. C., Calero-Nieto, F. J., Sánchez Castillo, M., Oedekoven, C. A., Diamanti, E., Schulte, R., Ponting, C. P., Voet, T., Caldas, C., Stingl, J., Green, A. R., Theis, F. J., & Göttgens, B. (2015). Combined Single-Cell Functional and Gene Expression Analysis Resolves Heterogeneity within Stem Cell Populations. Cell Stem Cell, 16(6), 712–724. 10.1016/j.stem.2015.04.004

61. World Malaria Report 2024. Addressing inequity in the global malaria response (2024). World Health Organization.

62. Yamamoto, R., Wilkinson, A. C., Ooehara, J., Lan, X., Lai, C. Y., Nakauchi, Y., Pritchard, J. K., & Nakauchi, H. (2018). Large-Scale Clonal Analysis Resolves Aging of the Mouse Hematopoietic Stem Cell Compartment. Cell Stem Cell, 22(4), 600–607.e4. 10.1016/j.stem.2018.03.013

63. Yamashita, M., Nitta, E., & Suda, T. (2015). Aspp1 Preserves Hematopoietic Stem Cell Pool Integrity and Prevents Malignant Transformation. Cell Stem Cell, 17(1), 23–34. 10.1016/j.stem.2015.05.013

64. Zioni, N., Bercovich, A. A., Chapal-Ilani, N., Bacharach, T., Rappoport, N., Solomon, A., Avraham, R., Kopitman, E., Porat, Z., Sacma, M., Hartmut, G., Scheller, M., Muller-Tidow, C., Lipka, D., Shlush, E., Minden, M., Kaushansky, N., & Shlush, L. I. (2023). Inflammatory signals from fatty bone marrow support DNMT3A driven clonal hematopoiesis. Nature Communications, 14(1). 10.1038/s41467-023-36906-1

